# A rapid, facile, and economical method for the isolation of ribosomes and translational machinery for structural and functional studies

**DOI:** 10.1101/2024.10.21.619433

**Authors:** Jessey Erath, Danielle Kemper, Elisha Mugo, Alex Jacoby, Elizabeth Valenzuela, Courtney F. Jungers, Wandy L. Beatty, Yaser Hashem, Marko Jovanovic, Sergej Djuranovic, Slavica Pavlovic Djuranovic

## Abstract

**Short abstract:** Ribosomes are essential RNA-protein complexes involved in protein synthesis and quality control. Traditional methods for ribosome isolation are labor intensive, expensive, and require substantial biological material. In contrast, our new method, RAPPL (RNA Affinity Purification using Poly-Lysine), offers a rapid, simple, and cost-effective alternative. This method enriches ribosomes and associated factors from various species and sample types, including cell lysates, whole cells, organs, and whole organisms, and is compatible with traditional isolation techniques. Here, we use RAPPL to facilitate the rapid isolation, functional screening, and structural analysis of ribosomes with associated factors. We demonstrate ribosome-associated resistance mechanisms from patient uropathogeni*c Escherichia coli* samples and generate a 2.7Å cryoEM structure of ribosomes from *Cryptococcus neoformans*. By significantly reducing the amount of the starting biological material and the time required for isolation, the RAPPL approach improves the study of ribosomal function, interactions, and antibiotic resistance, providing a versatile platform for academic, clinical, and industrial research on ribosomes.

**Long abstract:** Ribosomes are macromolecular RNA-protein complexes that constitute the central machinery responsible for protein synthesis and quality control in the cell. Ribosomes also serve as a hub for multiple non-ribosomal proteins and RNAs that control protein synthesis. However, the purification of ribosomes and associated factors for functional and structural studies requires a large amount of starting biological material and a tedious workflow. Current methods are challenging as they combine ultracentrifugation, the use of sucrose cushions or gradients, expensive equipment, and multiple hours to days of work. Here, we present a rapid, facile, and cost-effective method to isolate ribosomes from *in vivo* or *in vitro* samples for functional and structural studies using single-step enrichment on magnetic beads – RAPPL (RNA Affinity Purification using Poly-Lysine). Using mass spectrometry and western blot analyses, we show that poly-lysine coated beads incubated with *E. coli* and HEK-293 cell lysates enrich specifically for ribosomes and ribosome-associated factors. We demonstrate the ability of RAPPL to isolate ribosomes and translation-associated factors from limited material quantities, as well as a wide variety of biological samples: cell lysates, cells, organs, and whole organisms. Using RAPPL, we characterized and visualized the different effects of various drugs and translation inhibitors on protein synthesis. Our method is compatible with traditional ribosome isolation. It can be used to purify specific complexes from fractions of sucrose gradients or in tandem affinity purifications for ribosome-associated factors. Ribosomes isolated using RAPPL are functionally active and can be used for rapid screening and *in vitro* characterization of ribosome antibiotic resistance. Lastly, we demonstrate the structural applications of RAPPL by purifying and solving the 2.7Å cryo-EM structure of ribosomes from the *Cryptococcus neoformans*, an encapsulated yeast causing cryptococcosis. Ribosomes and translational machinery purified with this method are suitable for subsequent functional or structural analyses and provide a solid foundation for researchers to carry out further applications – academic, clinical, or industrial – on ribosomes.

---

As macromolecular RNA-protein complexes essential to the process of mRNA translation and quality control (*1–6*), ribosomes are highly conserved yet contain intrinsic diversity (*7–17*). The deconvolution of ribosome function and activity can provide insights into mechanisms of protein synthesis and quality control, as well as the state of a cell or organism at various times in its life cycle or differing growth conditions.

Methods have been developed to study ribosome-associated gene expression at the translation level - mainly ribosome purification by sucrose cushion or polysome profiling by sucrose gradients. Each method provided significant advancements to structural and functional studies of the mechanism of protein synthesis, gene expression control at the mRNA level, and quality control at the ribosome, mRNA, and nascent polypeptide levels. Although these methodologies offer valuable insights, as mentioned earlier, they are unsuitable for all material types, particularly those in low abundance. Purified ribosomes and translation-associated components end up in concentrated sucrose solutions, making downstream functional and structural applications cumbersome. Further purification and concentration steps usually result in additional loss of valuable material. Sucrose cushion and gradient purifications also require specialized equipment (ultracentrifuge, rotors, and fractionators) that can be cost-prohibitive for some labs and require significant time for proper separation. Due to lengthy procedures, these methods also require additional foresight and care regarding processes that could influence the chemical or biological stability of the material, oxidation status, and temperature control over long periods of time.

The interaction and coordination of cationic amino acids and anionic nucleic acid phosphate backbone have been well studied (*18*). On two occasions, monolithic anion exchange chromatography was used to isolate ribosomal subunits or ribosomes from partially purified or complex lysates of mycobacterium and baker’s yeast for downstream functional characterizations (*19, 20*). Though somewhat successful, these methods did not find a wider audience or everyday use. Homoamino acid polymers, particularly poly-lysine, have been widely used for various scientific applications. Its interactions and applications with nucleic acids have been documented for DNA extraction (*21*), DNA complex condensation (*22*), RNA purification (*23*), and to improve the delivery of siRNA complexes (*24*). However, poly-lysine has not been explicitly employed for the purification of cytoplasmic ribosomes (translating and non-translating), organelle-specific ribosomes (i.e. mitochondria, apicoplast or chloroplast), ribosomes undergoing biogenesis in the nucleus, and overall ribosome- or translation-associated factors. Moreover, the isolation of RNA using poly-lysine or similar moieties was limited to downstream applications. Such isolations were executed on denaturing RNA and protein molecules with complete disruption of the RNA-protein complexes. In such purifications, functional and structural information on RNA-protein complexes for further research was lost, and purified samples could not be used in both scientific and clinical applications for getting insight into ribosomes or translation machinery.

Here, we introduce RNA Affinity Purification by Poly-Lysine (RAPPL), a novel method for isolating a variety of functional RNAs. In this study, we focus on isolating ribosomes and translation-associated factors, with multiple examples where RAPPL was used to obtain structural and functional data on translational machinery. We demonstrate, quantitatively and qualitatively, significant enrichment of ribosomes from various biological samples, including some with limited material. The RAPPL-purified material can be used for multiple downstream applications such as mass spectrometry, imaging of translation drug effects, tandem purification assays, and *in vitro* translation assays. RAPPL can be combined with the previously described methods to enrich specific translation complexes and reduce material loss. Finally, to show the power of RAPPL, we perform a straightforward structural biology application of this new method in purifying ribosomes from the human intercellular pathogen *Cryptosporidium neoformans,* a difficult-to-process parasite. These ribosomes were structurally characterized by cryo-EM with a resolution of 2.7Å. Notably, prior to this work, *C. neoformans* ribosomes eluded structural studies because of their scarcity in purified samples. While RAPPL does not negate – and is compatible with – methods typically employed for this work, it provides a new means of obtaining RNA species for functional and structural study within the expanding translation field. This study focuses on only a subset of the potential applications of RAPPL, overcoming limitations of previous methods for ribosomes and associated translation machinery isolation. The method described here is affordable, straightforward, cost-efficient, and robust, allowing for the purification of high-quality ribosomes and related factors from different sample sizes and materials, from single-cell organisms and isolated/cultured cells to organs and whole organisms.

## Results

### Method Overview – RAPPL enriches ribosomes and associated factors

The full details of RAPPL are described in the Methods section. RAPPL is an anion exchange and affinity-based purification method that exploits the negatively charged backbone of RNA and positively charged ε-amino groups of poly-lysine at the physiologically relevant pH 7.5 (Figure 1A). For the purification of ribosomes and associated factors, cells or entire organisms are cultured and processed to specific application requirements. Lysis is performed in a typical ribosome isolation low-salt buffer containing DNase I, RNase inhibitor, and protease inhibitors. Crude lysate is clarified from cell membranes and non-soluble particles by short centrifugation and then bound to magnetic poly-lysine beads for an average of 15-30 minutes at 4. The beads are then washed in a buffer without detergent. Elution is carried out by incubating the beads in a wash buffer containing poly-D/L-glutamic acid for 15 minutes at either room temperature or 4. The procedure can be performed on an average time frame of 45-60 min, and the eluted material can be stored or applied to downstream applications. Compared with more traditional methods, the workflow is displayed in Figure 1A.

**Fig. 1:**
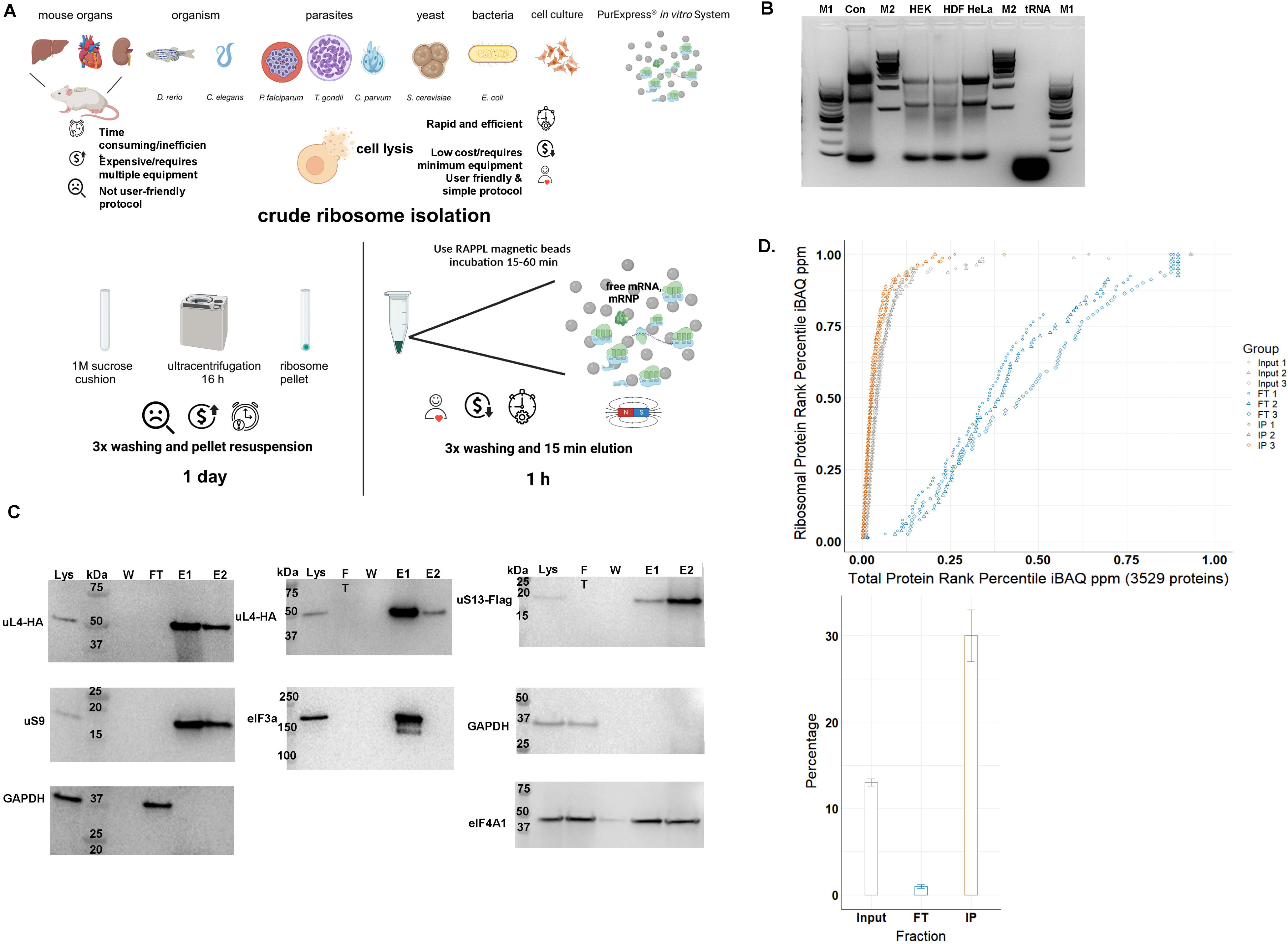
The RAPPL method. **A.** Schematic describing the advantages of RAPPL over conventional methods. **B.** 2% agarose gel of RAPPLE purified RNA samples from of HEK-293 (HEK), human dermal fibroblasts (HDF) and HeLa human cell cultures. RNA isolated from commercial HeLa cell lysate (Con; Thermo Fisher HeLa IVT kit) and purified yeast tRNAs (tRNA, Ambion) were loaded as controls. NEB 100 bp and 1 Kb base pair markers (M1 and M2, respectively) are used to estimate size of isolated RNAs. **C.** Western blot analysis of HEK-293 lines uL4-HA and uS13-Flag tagged by CRISPR/Cas9 throughout the RAPPL purification process – lysate (Lys), flow-through (FT), wash (W) and elution (E1 and E2). RAPPL is selective for ribosome-associated factors, showing that the HA-tagged ribosomal protein uL4, Flag-tagged uS13, as well as the untagged uS9 are in the elution fractions. Translation factors are also enriched and purified by RAPPL as seen by the visualization of eIF3A and eIF4A1 proteins by specific antibodies in elution fractions. Presence of GAPDH, as a control for loading is detected in lysate and flow-through. Molecular markers indicate size of detected proteins. **D.** (top) Plot of the of each HEK-293 ribosomal protein’s rank percentile in relation to total protein rank percentile for each replicate of input, flow-through (FT), and bead-bound (IP) fractions. (bottom) Graph representing percentage of ribosomal proteins in total protein associated with input, flow-through (FT), and poly-lysine bead-bound (IP) fractions. Error bars represent standard deviation of triplicate averages.

To determine the capacity and possibility of poly-lysine beads to bind ribosomes and translation-associated material, we incubated ribosomes from the PURExpress® *in vitro* translation system (NEB) with commercially obtained poly-lysine beads (Supplementary Figure 1). We visualized control and ribosome-bound beads via transmission electron microscopy (TEM). The ribosomes were visibly associated with magnetic particles coated with poly-lysine compared to beads without ribosome incubation (Supplementary Figure 1). The beads incubated with ribosomes also appear darker, with increased negative staining, and have less feathering at their periphery, suggesting dense ribosome binding over the surface of the beads. In support of this observation, adding elution buffer with poly-D-glutamic acid to the beads increased the number of ribosomes in imaging fields, arguing for strong surface binding of the ribosomes to the poly-lysine beads.

To assess the possible use of poly-lysine beads for the purification of ribosomes from complex lysates, we used lysates obtained from *E. coli* strain DH5α, HEK-293 cells, Human Dermal Fibroblasts (HDFs), and HeLa cells (Figure 1B and Supplementary Figure 2). After the RAPPL procedure (Figure 1A), we employed 20% of the total sample to determine the quality of the purified material. A typical yield from 5 to 10 million human cultured cells using 100 µL of slurry poly-lysine beads was 80-120ng. We used agarose gel electrophoresis for human cell lines (Figure 1B). All three RAPPL purified samples from human cell lines indicated significant amounts of 28S and 18S rRNAs and tRNAs based on controls, commercial *in vitro* translation lysate from HeLa cells, and purified yeast tRNAs, respectively. *E. coli* DH5α isolated ribosomes were analyzed by bioanalyzer (Supplementary Figure 2). A typical RAPPL total RNA yield from 50 mL *E. coli* cell culture lysate was 100-120ng. The analyzed sample indicated significant enrichment of both 16S and 23S rRNAs and shorter RNA species.

To further demonstrate RAPPL enrichment of ribosomes and translation-associated factors, such as ribosomal proteins and initiation factors, over those not associated with protein synthesis, we turned to CRISPR/Cas9-engineered HEK-293 cell lines. RAPPL was performed using lysates from HEK-293 cell lines in which uS13 (RPS18) and uL4 (RPL4) have been Flag- and HA-tagged, respectively, by insertion of tag sequences in endogenous loci of the appropriate gene (Supplementary Figure 3). Western blot analyses of RAPPL purified HEK-293 samples indicated complete binding of tagged ribosomal proteins (uS13, uL4) to poly-lysine beads without a visible band in either flow-through or wash fractions (Figure 1C). In addition to the tagged proteins, we have tested the binding of non-tagged 40S subunit ribosomal protein uS9 (RPS16), eIF3a, and eIF4A1 initiation factors, and GAPDH protein. While uS9 and eIF3a were readily detectable in poly-lysine bound fractions, heavily abundant eIF4A1 was detectable in all fractions, whereas GAPDH remained only in the flow-through fraction during the RAPPL procedure (Figure 1C).

Finally, to fully detail the repertoire of proteins enriched by RAPPL, the purification procedure was performed on HEK-293 cells and *E. coli* strain DH5α lysates in triplicate, followed by quantitative mass spectrometry analysis (Figure 1D, Supplementary Figure 4, and Supplementary Table 1 and 2). We directly compared ribosomal proteins to all other proteins by abundance in cell lysate (input), flowthrough (FT), and poly-lysine bead-bound (IP). We observed enrichment in ribosomal proteins in poly-lysine bound fractions from *E. coli* and HEK-293 cells (Figures 1D and Supplementary Figure 4).70% and 30% of all poly-lysine bound proteins were ribosomal proteins in the case of *E. coli* and HEK-293 cell pull-downs, respectively. We detected 81 annotated human ribosomal proteins in HEK-293 pull-down (Supplementary Table 1) and 52 annotated E. coli ribosomal proteins (Supplementary Table 2). Moreover, after incubation with poly-lysine beads, the flowthrough fraction of HEK-293 cell lysate was almost entirely depleted from ribosomal proteins (Figure 1D). Human ribosomal proteins represented 1% of total proteins in flow-through fraction, compared to approximately 13% in starting lysate and 30% of poly-lysine bound fraction (Figure 1D). These results further confirmed ribosome enrichment and efficient binding to poly-lysine beads seen previously in western blot analysis of multiple ribosomal proteins (Figure 1C). In addition to the enrichment of ribosomes, as seen by the analysis of ribosomal proteins, we also noticed enrichment in ribosome- and translation-associated proteins (Supplementary Tables 1 and 2). In the top 100 proteins from *E. coli* bound to poly-lysine beads were translation initiation factor 3 (*infC*), elongation factor Ef-Tu (*tufA, tufB)*, rRNA processing and maturation factors (*rbfA, hpf, raiA, rnr, rimM*), along with ribosome and nascent polypeptide chain associated proteins such as trigger factor (*tig*) and chaperons (*groL*) (Supplementary Table 2). In the poly-lysine bound data from HEK-293 cells, we could readily detect enrichment of nascent polypeptide chain associated chaperons (HSP70), translation elongation factors (EEF1 and EEF2), all 13 members of eIF3 translation initiation complex, eIF4A1, eIF5 as well as eIF6, among others (Supplementary Table 1). Notably, enrichment of eIF3A and eIF4A1 proteins was also previously detected using western blot analyses of poly-lysine bound fractions (Figure 1C).

As such, by exploiting the negatively charged RNA backbone and the positively charged ε-amino groups of poly-lysine, we can isolate ribosomes of high quality with a relatively fast protocol from cell lysate. Using RAPPL, we isolate not only ribosomes but most of the expected translation-associated factors (IF-1, IF-2, IF-3, EF-Tu, EF-Ts, EFG EFP, among others in *E. coli* samples, as well as all 13 members of eIF3 complex, eIF4A, eIF5, eIF6, EEF1, EEF2, among others in HEK293 samples). We also isolate many factors that have been suspected to be associated with ribosome and translation (i.e., YhbY (*25*) and YibL (*26*) in *E. coli*, or LARP1 (*27*) and SERBP1 (*28*) in HEK293 cells), as well as multiple new candidates that need further confirmation.

### RAPPL overcomes material scarcity limitations

Sample limitations present a major challenge for many purification processes, including those used for ribosomes and translation-associated material. This is often the case for clinically relevant samples, specific cell types, or organs, and it is often exaggerated in parasitology, with intercellular parasites being present in small numbers and at certain stages of various parasite life cycles. We, therefore, tested the lower limits of RAPPL by performing purifications on decreasing ribosome or cell numbers (Figure 2). We first used ribosomes from the PURExpress® System, which provides purified and highly concentrated *E. coli* ribosomes in a known quantity (13.3 mM). We diluted *E. coli* ribosomes in RAPPL lysis buffer starting at 13.3 μM down to 1.3 nM. The ribosomes were then purified using RAPPL, and the eluates were examined by TEM. We could show ribosome isolation from the lowest concentration at 1.3 nM (Figure 2A).

**Fig. 2:**
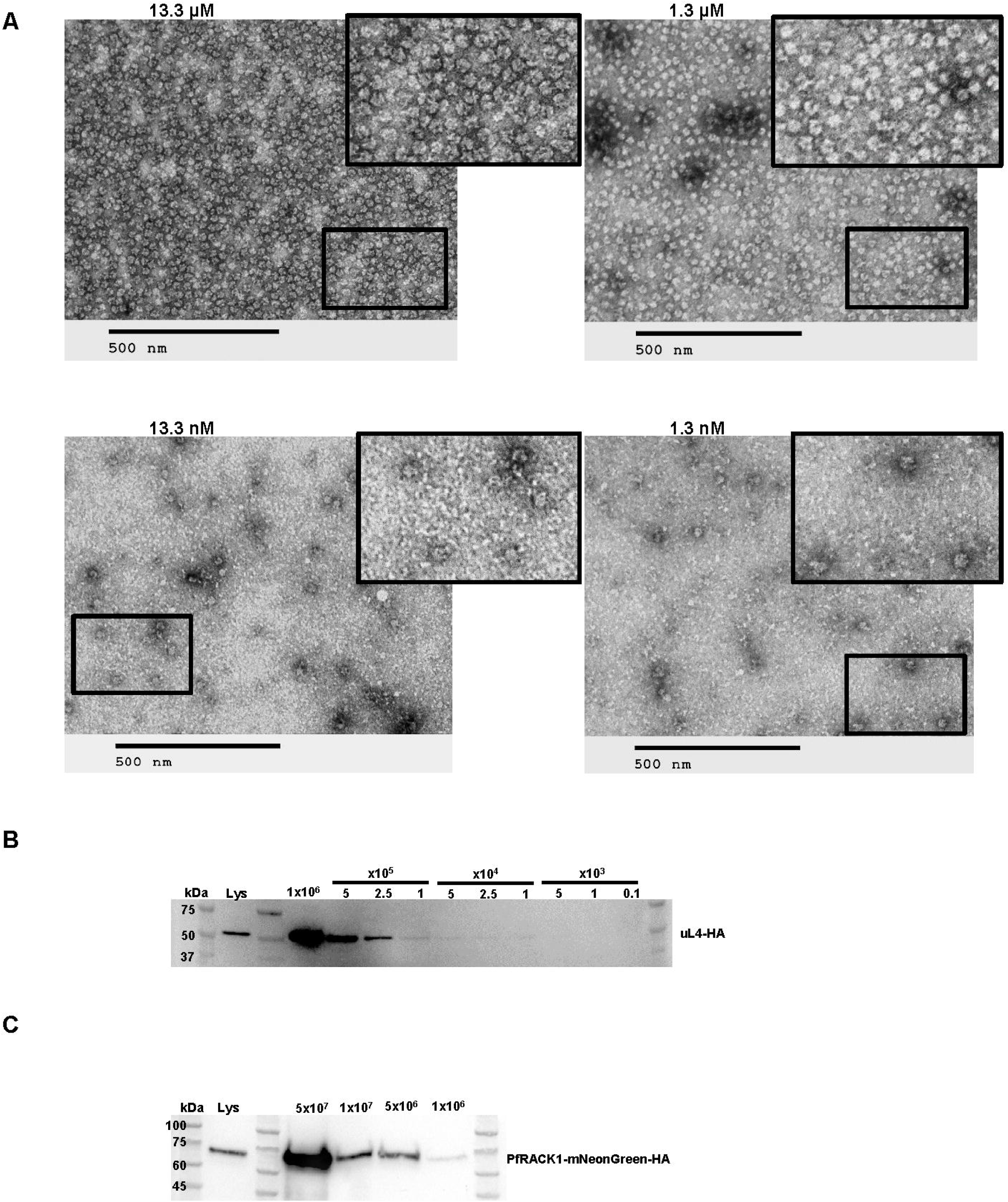
RAPPL can enrich and purify ribosomes from limited biological material. **A.** TEM visualization of RAPPL eluates from PureExpress® ribosomes from a 10-fold dilution scheme of 13.3 μM to 1.3 nM. The scale bar represents 500 nm. **B.** Western blot analysis using αHA antibody on RAPPL eluates of HEK-293 cells in which uL4 was HA-tagged by CRISPR/Cas9 whereby the starting cells were diluted to 1×10^6^, 5×10^5^, 2.5×10^5^, 1×10^5^, 5×10^4^, 2.5×10^4^, 1×10^4^, 5×10^3^, 1×10^3^, and 0.1×10^3^ cells prior to lysis. **C.** Western blot analysis using αHA antibody on RAPPL eluates of *P. falciparum* NF54 cells in which PfRACK was C-terminally tagged with mNeonGreen-HA whereby the starting cells were diluted to 5×10^7^, 1×10^7^, 5×10^6^, and 1×10^6^ cells. Molecular markers indicate size of detected proteins.

To examine method limitations in the context of cell lysates, components of which could easily affect ribosome binding, we performed RAPPL on HEK-293 cells where uL4 (RPL4) and uS4 (RPS9) have been HA- or Flag-tagged by CRISPR/Cas9 engineering, respectively (Supplementary Figure 3). We performed a series of cell dilutions, and the lysates were used further for RAPPL. The eluates of RAPPL isolation were then analyzed by western blot using αHA-HRP or αFLAG antibody to detect the tagged uL4 (Figure 2B) or uS4 (Supplementary Figure 5) proteins, respectively. Results indicated a lower detection limit by western blot at 5,000 or 10,000 cells for uL4 and uS4 tagged cell lines, respectively (Figure 2B and Supplementary Figure 5).

Mammalian cell lines can be easily cultured in high abundance, with larger cells containing significantly more ribosomes than other clinically significant organisms, such as the malaria-causing parasite - *Plasmodium falciparum*. Growing and maintaining synchrony of the large parasite cultures in replicates necessary for some studies is difficult and costly. Additionally, growing parasites to high parasitemia to reduce flask numbers and materials generates stress conditions that confound results and isolation of ribosomes. We, therefore, wanted to test the cell number limitations of RAPPL to purify from *P. falciparum* NF54 cell line. We engineered this *P. falciparum* cell line with CRISPR/Cas-9 by inserting an HA-tagged mNeonGreen reporter in the C-terminus of ribosomal RACK1 protein (*Pf*RACK1-mNeonGreen-HA, Supplementary Figure 6). Parasites were synchronized at the ring-stage, grown to ∼5% parasitemia in 3% hematocrit. Late-stage parasites were isolated via MACS magnet purification (*29*). Parasites were counted using the countess cell counter. Cells were then diluted to 5×10^7^, 1×10^7^, 5×10^6^, and 1×10^6^ cells, lysed, and the clarified lysate used in the RAPPL method. The products were analyzed by western blotting with αHA-HRP or α-uS11 (RPS14) antibody (Figure 2C and Supplementary Figure 7). Our results indicate that we can detect tagged or non-tagged ribosomal proteins isolated from down to one million *P. falciparum* cells. Our results indicate that RAPPL can purify ribosomes and translation-associated factors from relatively small quantities of starting material. Current detection levels associated with 1 nM *E. coli* ribosome concentration and western blot analysis of 5000 RAPPL purified HEK-293 or 1×10^6^ of *P. falciparum* cells.

### RAPPL is amenable to a wide variety of material and organism types

To test whether the RAPPL method may be used as a versatile tool for isolating ribosomes regardless of starting material, we applied RAPPL to a range of sample types, from single-cell organisms and cultured cells to tissues and whole organisms. In each case, we isolated translation-associated materials (i.e., ribosomes) that we then visualized by TEM (Figure 3). RAPPL was efficient in isolating ribosomes regardless of starting material type and quantity. However, slight modifications to the lysis step were necessary but amenable to ensure success for each sample. To ensure the lysis of single-cell organisms (Figure 3A) (i.e., *E. coli*, *S. cerevisiae*, *T. gondii*, *P. falciparum*, and *C. parvum*), cell breaking was done by bead-beating. For samples containing high tissue organization (i.e., perfused mouse organs and *C. elegans*), they were flash frozen, resuspended in buffer, and bead-beating was used to lyse them (Figure 3B). The *D. rerio* was further processed by first finely scoring and segmenting the specimen using a scalpel, followed by flash-freezing in liquid nitrogen and pulverization using a mixer miller (Figure 3B). Lysis of all samples was performed in similar buffers with two exceptions. In the case of *P. falciparum* cells, we used well-documented specific lysis buffer conditions necessary for ribosome isolation and stripping of ribosomes from the endoplasmic reticulum (*30*). This was also applied to *C. parvum* sporozoites. Additionally, high levels of heme present in red blood cells were avoided by using the previously mentioned MACS magnet purification of the parasite (*29*). RAPPL is, therefore, highly adaptable to various starting materials.

**Fig. 3:**
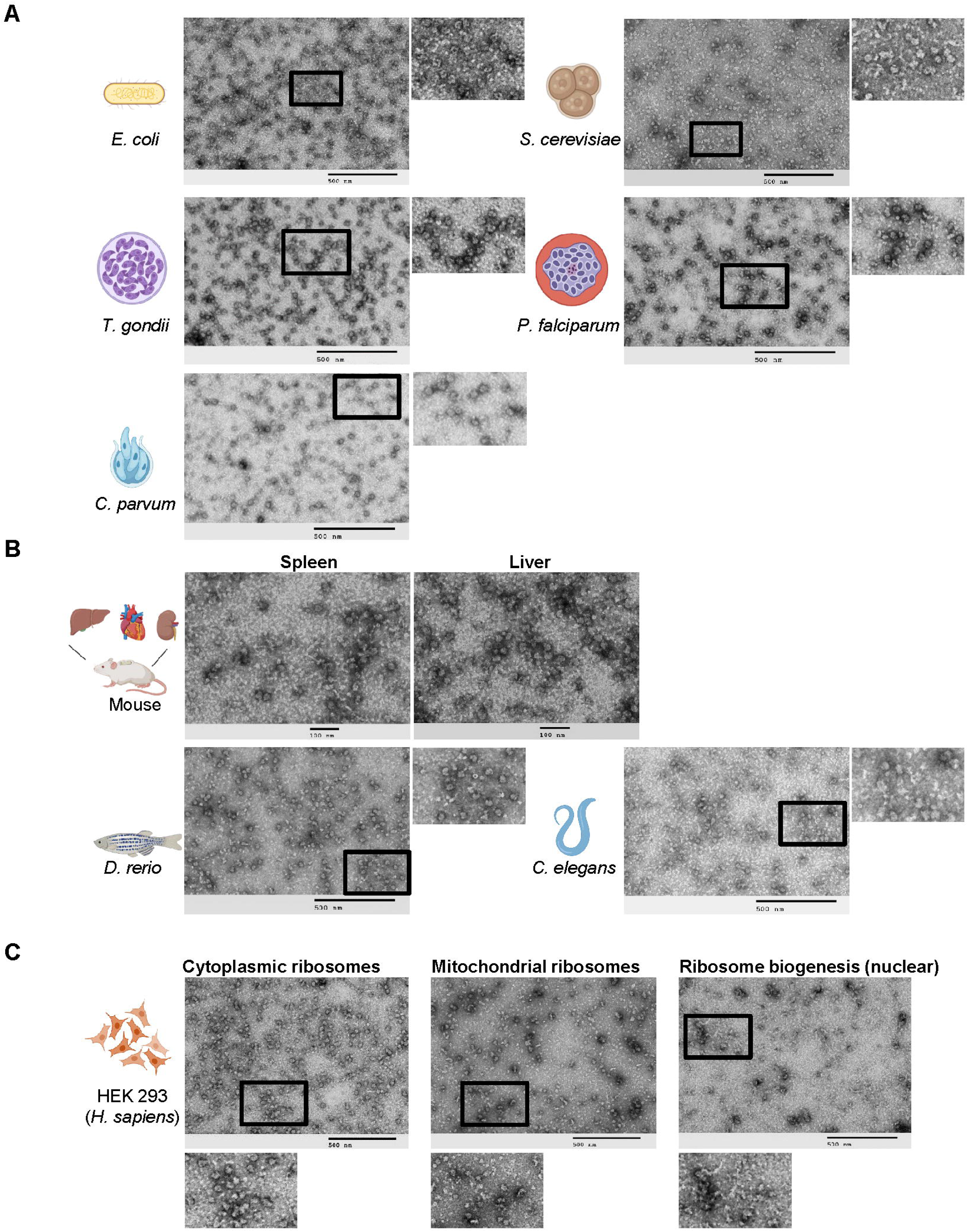
RAPPL is a versatile method multiple single celled organisms, tissues, and multicellular organisms. **A.** TEM visualization of RAPPL eluates purified from several single celled organisms: *E. coli*, *S. cerevisiae*, *T. gondii*, *P. falciparum*, *C. parvum*. **B.** TEM visualization of RAPPL eluates purified from mouse tissue sections of spleen and liver as well as whole organisms *D. rerio* and *C. elegans*. **C.** TEM visualization of compartment-specific ribosomes generated from RAPPL eluates of cytoplasmic, mitochondrial, and nuclear (ribosome biogenesis) fractions. The scale bar represents 100 or 500 nm.

In contrast to bacterial cells, eukaryotic cells contain organelle-specific ribosomes. Mass-spectrometry analyses indicated that RAPPL enriches both cytoplasmic and mitochondrial ribosomes (Figure 1D, Supplementary Figure 4, and Supplementary Table 1-3). Besides subcellular compartment-localized ribosomes (e.g., mitochondria and chloroplasts in plants), ribosome biogenesis is compartmentalized in the nucleus. Mass spectrometry analyses of HEK-293 lysate also indicated enrichment of multiple ribosome biogenesis factors using RAPPL (Supplementary Table 2). As such, to further indicate the versatility of our method, we sought to separate compartment-localized ribosomes from cytoplasmic ones (Figure 3C). We isolated the mitochondrial, nuclear, and cytoplasmic cell fractions using the previously published method (31) or commercially available kits (32–34). Separation of cytoplasmic and mitochondrial fractions was done in CRISPR/Cas9 engineered uS4-Flag tag HEK-293 cell line while separation of cytoplasmic and nuclear fractions was executed in uL4-HA-tagged HEK-293 cell line. Lysates from cellular compartments and cytoplasm were used as a starting point for RAPPL procedures.

The RAPPL eluates were then visualized by TEM (Figure 3C). Images of each fraction contained unique sets of ribosome-like particles indicating the separation of cytoplasmic-, mitochondrial-, and nuclear-associated ribosome particles. We used western blot analyses to validate the separation of cellular compartments and enrichment of ribosomal particles (Supplementary Figure 8). To demonstrate cytoplasmic and mitochondrial fraction separation, we used a Flag antibody for uS4-Flag and mitochondrial ribosomal protein S35 (mRPS35) specific antibody for cytoplasmic and mitochondrial ribosomes, respectively. uS4-Flag protein was strongly enriched in cytoplasmic lysate and RAPPL eluate, while mRPS35 was only detected in mitochondrial lysate and RAPPL eluate (Supplementary Figure 8A). In the case of nuclear and cytoplasmic fractions, we used HA antibody to detect the uL4-HA protein. Analysis revealed the presence and enrichment of large ribosomal subunit uL4 protein in both cytoplasmic and nuclear lysates and RAPPL eluates (Supplementary Figure 8B). The ratio of uL4 in lysate and RAPPL elution followed previously observed enrichment in RAPPL eluates, indicating that most of the uL4 protein is present in the cytoplasmic fraction of HEK-293 cells. A small portion of uL4 ribosomal protein was detected in the nucleus, possibly involved in ribosome biogenesis. Topoisomerase II β-specific antibody was used to confirm the separation of nuclear fraction (Supplementary Figure 8B). Taken together, these results indicate that we can fractionate cell compartments and use RAPPL to isolate cytoplasmic, nuclear, and mitochondrial ribosomal particles to obtain insight into compartment-specific translation machinery in the case of the mitochondria or into ribosomes undergoing biogenesis from the nuclear fractions. These results demonstrate that RAPPL can robustly isolate translation-associated material from a wide range of substances – single cells, tissues, or whole organisms. The method is further adaptable to the ribosome isolation requirements of different organisms or cellular compartments (cytoplasmic, mitochondrial, nuclear, among others), making it a versatile new tool.

### RAPPL is compatible with current technologies and methodologies for the study of protein synthesis

To further test the applicability of the RAPPL method, we sought to determine whether the ribosomes and associated translation factors generated by RAPPL can be used to study protein synthesis, the effects of known translation-associated drugs, or the purification of translation-associated complexes. We first used commercial eukaryotic *in vitro* translation kits as proof of concept (Supplementary Figure 9). We directly bound wheat germ and HeLa cell lysates to the RAPPL beads or mixed lysates with eGFP mRNAs, and after protein synthesis was carried on for 2 hours, samples were subjected to RAPPL. The eluates were then visualized by TEM. The micrographs show that the translation architecture is maintained throughout the purification, whereby ribosomes remain bound to mRNA and indicate possible organization in polysomes (Supplementary Figure 9).

To determine if RAPPL can be used to assay the effects of different drugs on cultured cells, we performed RAPPL using HEK-293 cells treated and lysed in the presence of various translation inhibitors. Visualization by TEM showed inhibitor-dependent variation in ribosome organization versus untreated controls (Figure 4A). The organization of ribosomes treated with translation elongation inhibitors cycloheximide and anisomycin tend to show more polysomes (i.e., ‘beads on a string’). In contrast, the translation initiation inhibitor harringtonine reduced this effect by having more monosomes (i.e., individual ribosomes). Control samples (without inhibitors) showed the combination of monosomes, some disomes, and separated subunits (Fig 4A). Naturally, cryo-EM at high resolution can confirm the observations mentioned above beyond any doubt. However, the idea behind the presented panels of Figure 4A is to show that the incubation with poly-lysine beads and the subsequent elution do not seem to interfere with the global translation landscape.

**Fig. 4:**
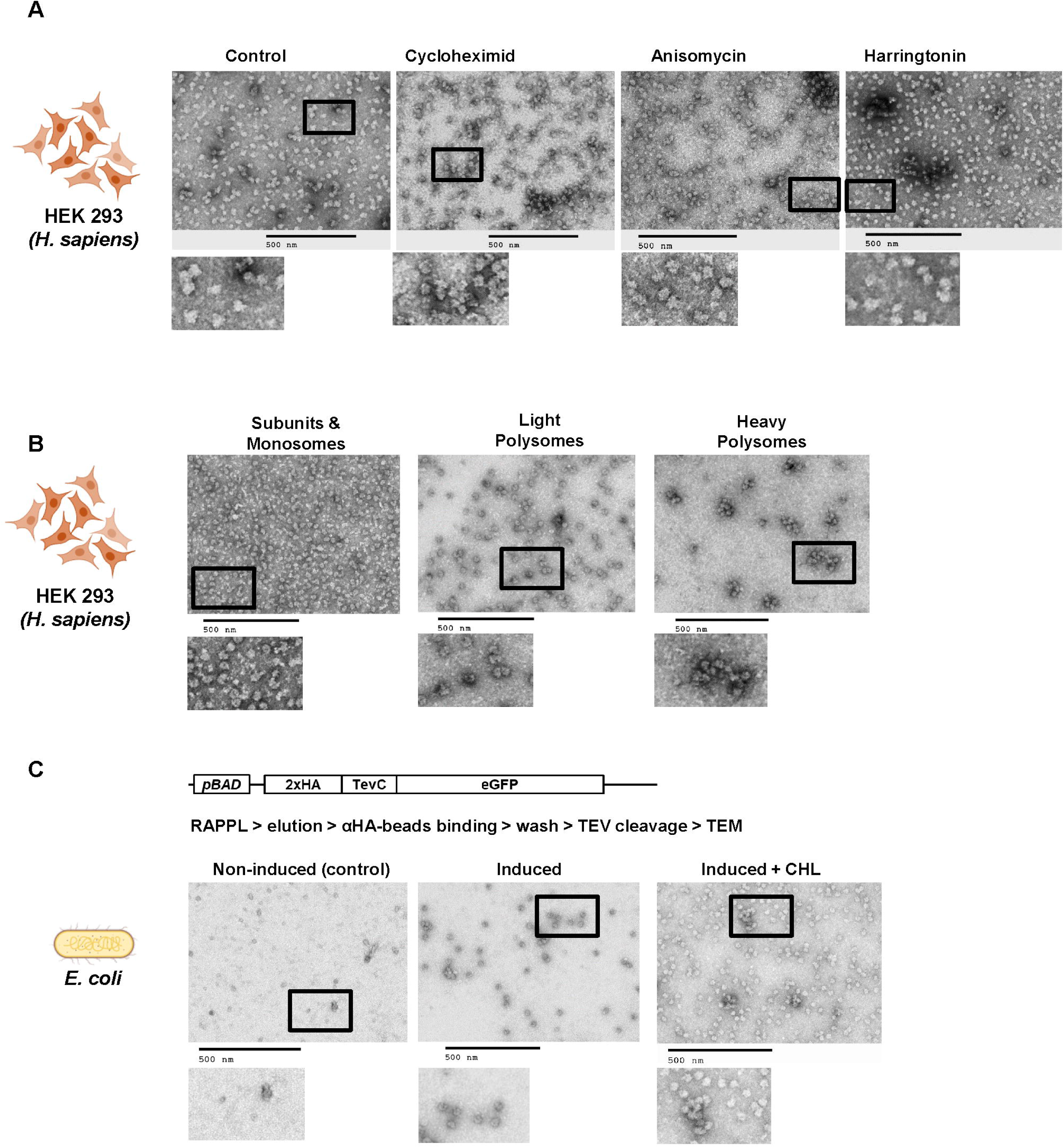
The elutions of RAPPL can be used in downstream applications. **A.** HEK-293 lysates were treated with cycloheximide, anisomycine, and harringtonine with untreated lysate as a control. Ribosomes were purified using RAPPL in the presence of inhibitors and the eluates visualized by TEM. **B.** HEK-293 lysates were fractionated using polysome profiling. Fractions corresponding to ribosome subunit and monosomes, light polysomes, and heavy polysomes were pooled, respectively. These pools were diluted 1:5 to ensure that sucrose did not interfere with binding. The diluted, pooled samples were subject to RAPPL and the eluates visualized by TEM. **C.** Schematic of arabinose-inducible reporter expressing a 2xHA affinity tagged eGFP reporter separated by a TEV protease cleavage site (top). RAPPL was performed on bacterial lysates in the absence or presence of bacterial translation elongation inhibitor chloramphenicol (CHL) followed by αHA magnetic bead purification, again ±CHL, finally eluting with TEV protease. Eluates were visualized by TEM. Non-induced are shown as controls for lack of protein production and subsequent non-specific binding to αHA beads. The scale bar represents 500 nm.

Polysome profiling is an invaluable tool for studying many aspects of protein synthesis. However, isolating RNA and protein from the generated fractions is cumbersome, with a significant sample loss (*20*). To determine if RAPPL was compatible with these sucrose-containing fractions, we performed polysome profiling using HEK-293 cells. The fractions for the subunits/monosomes, light polysomes, and heavy polysomes were pooled, respectively, and subject to RAPPL. The eluates were then visualized by TEM (Figure 4B). Our results show that RAPPL is compatible with polysome profiling, and ribosome organization is once again maintained (Figure 4B), allowing our method to enrich ribosomes from these fractions while removing the contaminating sucrose for further analysis.

Finally, we wanted to know whether RAPPL can be used as a starting point for further purification of certain translation complexes. Tandem affinity purifications often isolate specific translation complexes or improve the final sample purity. Therefore, we performed a tandem RAPPL-αHA-bead purification using *E. coli* cells in which the eGFP construct, N-terminally tagged with a double HA tag followed by an engineered TEV-protease cleavage site, was expressed under an arabinose inducible promoter (Figure 4C). In this case, we wanted to isolate translation complexes associated with the nascent polypeptide chain of the eGFP reporter. The expression of eGFP was induced by the addition of arabinose in the media. Non-induced cells served as a control. Cells were lysed in the absence or presence of chloramphenicol (CHL) - a translation elongation inhibitor for *E. coli* ribosomes. The clarified lysates were subjected to RAPPL and eluates from poly-lysine beads were then incubated with αHA magnetic beads. The αHA beads were then washed and eluted with the addition of His-TEV Protease in the wash buffer. The αHA-bead eluates were then analyzed by TEM (Figure 4C). Images of control non-induced samples did not contain any ribosomes, while induced samples indicated the presence of ribosomes or even polysomes. The observed difference in the number of ribosomes and observed polysomes in induced samples was attributed to CHL. Addition of CHL during lysis prevents run-off of elongating ribosomes and release of the nascent polypeptide chain, resulting in a higher number of ribosomes associated with HA-beads and in RAPPL eluate from CHL-treated lysates (Figure 4C).

Therefore, RAPPL can be used with current technologies and methods to study protein synthesis. RAPPL allows for sample enrichment from and removal of sucrose, which is often incompatible with downstream techniques. Ribosome binding and organization is maintained throughout the purification process, suggesting efficacy in structural applications and the possibility to use RAPPL as an enrichment step for tandem purification of specific translation complexes with tagged target proteins. Finally, translation inhibitors may be used to perturb ribosomes and translation factor equilibrium in such studies to obtain the desired fractions and purity of complexes of interest.

### RAPPL eluates are compatible with functional and downstream clinical applications

While the products of RAPPL appear to have the visual hallmarks of functional translation, we wanted to determine if RAPPL-isolated ribosomes maintain their functionality – essentially, could RAPPL-isolated ribosomes translate reporter mRNA into protein. To test the activity of isolated ribosomes, we used *E. coli* cells and a widely used PURExpress® *in vitro* translation system kit (*31*). We first tested whether adding poly-D-glutamate, used for ribosome elution in the RAPPL method, would affect PURExpress® *in vitro* translation of an eGFP reporter. We did not observe any difference in the yield of eGFP protein synthesized by the PURExpress® kit with or without adding poly-D-glutamate (Supplementary Figure 10A). We next substituted the kit-supplied ribosomes with an increasing amount of RAPPL eluate from *E. coli* DH5α cells mixed with a DNA template encoding the eGFP reporter gene (Figure 5A). The estimated concentration of RAPPL-isolated ribosomes used in *in vitro* translation was 20-100 times lower than those used in the PURExpress® kit (approximately 2.4 μM). The reaction products were analyzed using a western blot to detect eGFP protein (Figure 5A). Results demonstrated the synthesis of eGFP reporter that is dependent upon the addition of increasing amounts of RAPPL-isolated ribosomes, and as such indicated that performing RAPPL on *E. coli* DH5α cell lysates resulted in enrichment of functional ribosomes, which can be used for *in vitro* translation systems. The control reaction (without a DNA template) displayed no eGFP synthesis (Figure 5A). We were also able to follow eGFP reporter protein synthesis from RAPPL-isolated ribosomes by following the fluorescence of newly synthesized reporter protein using a plate reader (Figure 5B). This further corroborates the robustness and functionality of RAPPL-isolated ribosomes and creates a simple assay to follow the functionality of RAPPL-isolated ribosomes. In addition to RAPPL purified and eluted ribosomes used in the solution mentioned above assays (Figure 5A and 5B), incubation of RAPPL beads used to isolate *E. coli* DH5α ribosomes with the PURExpress® buffer and DNA template resulted in active translation and synthesis of eGFP protein analyzed by western blot analyses (Supplementary Figure 10B). As such, RAPPL ribosome eluates or on-bead isolated ribosomes could perform *in vitro* translation reactions and translate reporter genes into protein, drastically shortening *in vitro* translation assays (Figure 5A,5B and Supplementary Figure 10B).

**Fig. 5:**
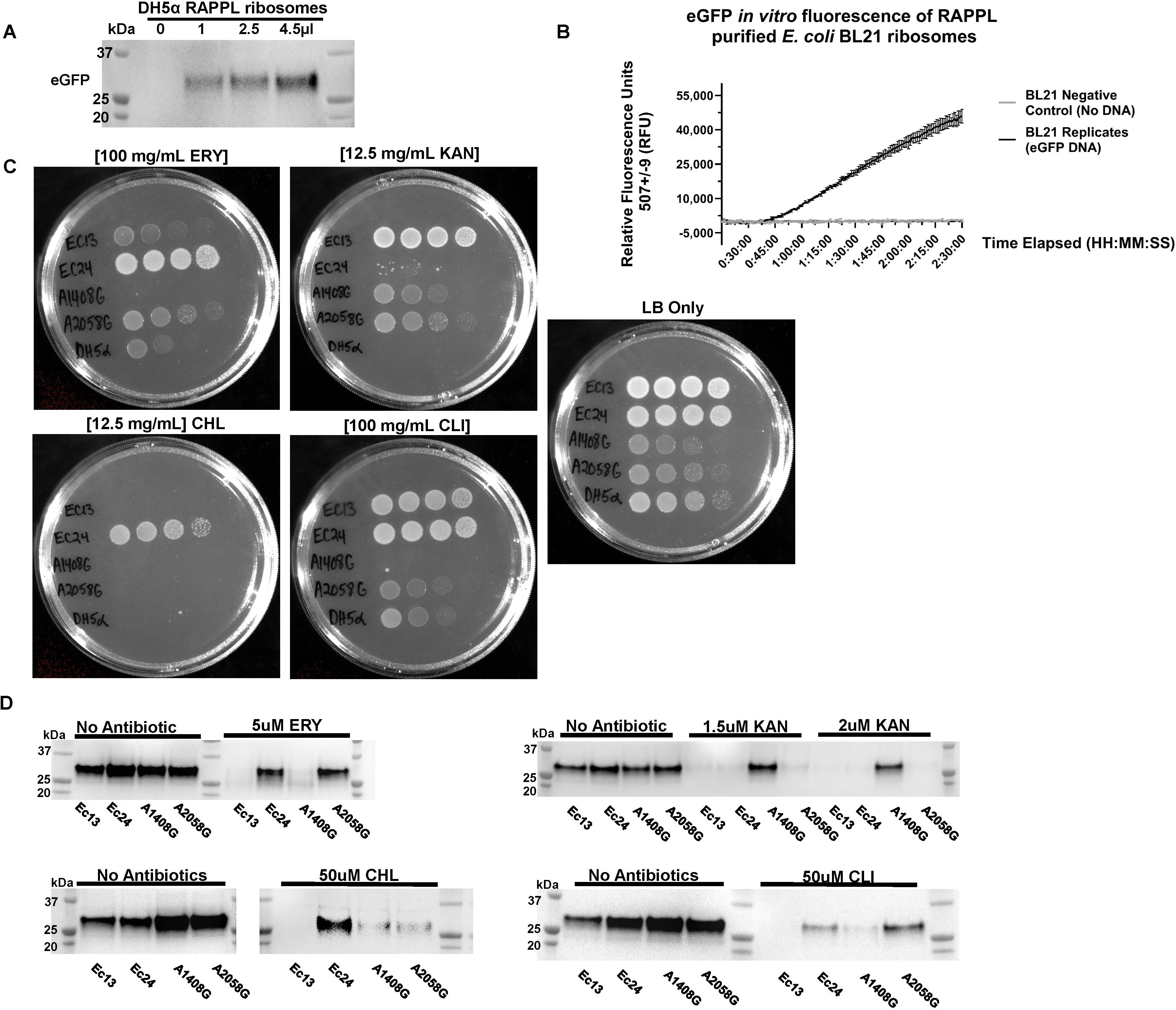
RAPPL isolated ribosomes are translationally active and can be used for clinical applications. **A.** Ribosomes were purified by RAPPL from *E. coli* DH5α cells grown to exponential phase, eluting in 30 μL of RAPPL elution buffer. Eluates were then used in the PURExpress® *in vitro* translation system instead of kit ribosomes. A PCR product encoding for eGFP harboring the T7 promoter and a polyA tail was used in the reaction (See Method for full details). Reactions were incubated for four hours. Ribosomes purified using RAPPL are active and able to translation mRNA. **B.** Activity of RAPPL purified E. coli BL21 ribosomes in the in vitro PURExpress® assays were observed using a kinetics protocol measuring eGFP fluorescence on an imaging plate reader. Relative fluorescence was determined with excitation settings set to wavelengths of 488 ± 9 and emission settings set to wavelengths of 507 ± 9. The standard deviation of technical triplicates eGFP fluorescence activity over a two hours and 30 minutes period with and without the DNA template encoding for eGFP are shown on the graph. **C.** Plate bacterial growth assays were performed using erythromycin (ERY), kanamycin (KAN), chloramphenicol (CHL), and clindamycin (CLI) to demonstrate strain resistance with LB only as controls for growth and the DH5α strain was used as a control strain. Concentration of used antibiotics is indicated. **D.** Synthesis of eGFP reporter by RAPPL isolated ribosomes in the absence and presence of indicated antibiotics (ERY, KAN, CHL, and CLI) targeting *E. coli* ribosomes. For each strain, 4.5 μL of 1.5 µg / µL of RAPPL isolated ribosomes was used for standard 25 μL PURExpress® *in vitro* Δ ribosome translation reaction (See Method for full details). Western blot analysis was performed on samples collected after 4 hours of incubation at 37°C and visualized using αGFP specific antibody. Molecular markers indicate size of eGFP protein.

To further test whether fast isolation of ribosomes by RAPPL could be used in clinical applications, we applied our method to test antibiotic-resistant *E. coli* strains associated with urinary tract infections. We used previously described patient clinical isolates of uropathogenic *E. coli* (UPEC) - Ec13 and Ec24 (*32*). These *E. coli* strains are trimethoprim-sulfamethoxazole and ciprofloxacin-resistant and have a broad-spectrum secondary multidrug transporter (MdfA+), providing additional antibiotic resistance (*33*). The Ec24 strain has been confirmed to contain active rRNA methylase (ermB), which results in the methylation of 23S rRNA (A2058), reducing erythromycin (ERY) binding to ribosomes (*34*). Additional E. *coli* strains with plasmid-encoded rRNAs and engineered rRNA mutations were also used as controls (*35*). The SQ110 L\TC 16S – A1408G strain carries a mutation that provides resistance to spectinomycin, kanamycin (KAN), and gentamicin, while the SQ110 L\TC 23S – A2058G provides resistance to spectinomycin and ERY. The SQ110 plasmid carries a selection cassette that encodes aminoglycoside 3’-phosphotransferase II enzyme (*35*) that inactivates KAN by phosphorylation in cells. However, ribosomes isolated from L\TC 23S – A2058G strain should not have resistance to KAN. Although both strains are kanamycin resistant and can grow on KAN-containing bacterial agar plates, only a small subunit rRNA mutation (A1408G) provides ribosome resistance to KAN (*35*). The SQ110 ΔTC 23S – A2058G plasmid contains a mutation in the large subunit rRNA (A2058G) that provides resistance to ERY and CLI (*35*). Using *E. coli* lab strain DH5α as a control, we tested antibiotic resistance for each *E. coli* strain by growing them on bacterial agar plates without (LB only) and supplemented with antibiotics (ERY, KAN, chloramphenicol (CHL), and clindamycin (CLI); (Figure 5C). All *E. coli* strains were able to grow on a plate without antibiotics (LB only). SQ110 strains grew slower than other *E. coli* strains due to the fact that all ribosomes in these strains are encoded by a single copy of rDNA located on a plasmid. Based on agar plate growth, the Ec13 UPEC strain was not resistant to CHL but to KAN, CLI, and partially to ERY (Figure 5C). The Ec24 UPEC strain was resistant to ERY, CLI, and CHL, while no resistance to KAN was observed (Figure 5C). As expected, SQ110 strains were resistant to KAN due to the plasmid antibiotic selection cassette and A1408G mutation in the 16S rRNA of SQ110 L\TC 16S – A1408G strain (*35*)(Figure 5C). SQ110 L\TC 16S – A1408G did not show resistance to any other antibiotics. Besides plasmid-encoded KAN resistance, SQ110 L\TC 23S – A2058G also indicated no resistance to CHL and strong resistance to ERY and CLI, due to engineered A2058G mutation in 23S rRNA (Figure 5C). *E. coli* DH5α showed partial antibiotic resistance to ERY and CLI and no resistance towards KAN or CHL (Figure 5C). The partial resistance of *E. coli* DH5α on ERY and CLI plates is due to 100 mg/mL of ERY and CLI used for agar plates.

To test whether observed antibiotic resistance is due to the specific methylation or mutation of ribosome nucleotides versus multidrug transporters, we used the RAPPL method on cell lysates from small bacterial cultures (volume of 50 mL). Each *E. coli* strain was grown to an exponential phase, cells were harvested, and RAPPL was performed. The RAPPL eluates from each strain were then used in the PURExpress® *in vitro* translation system to synthesize the eGFP reporter, as shown above (Figure 5A). All RAPPL purified ribosomes were tested for functionality in the control conditions (con, no antibiotic added in *in vitro* translation reaction; Figure 5D). After successful testing and indication of robust synthesis of eGFP reporter by western blot analyses, we carried on by testing the same set of RAPPL-isolated ribosomes but with the addition of antibiotics, previously used for testing of growth on agar plates (Figure 5C). We used antibiotic concentrations based on previous studies with bacterial resistance strains and engineered ribosome mutations (*35–38*). Only Ec24 and SQ110 L\TC 23S – A2058G RAPPL-isolated ribosomes were able to *in vitro* synthesize eGFP protein in the presence of 5 μM ERY or 50 μM CLI (Figure 5D), thus confirming the presence of the ermB methylase in the Ec24 strain and engineered A2058G mutation in the L\TC 23S – A2058G strain, respectively. In addition to ERY and CLI resistance, Ec24 RAPPL-isolated ribosomes indicated functional resistance in the presence of 50 μM CHL in an *in vitro* translation reaction (Figure 5D), confirming previous growth of this *E. coli* strain on agarose plates supplemented with CHL (Figure 5C). We did not observe any other RAPPL-isolated ribosomes with CHL resistance. Ribosomes isolated from SQ110 L\TC 23S – A2058G, as well as Ec24 ribosomes, could not synthesize eGFP in *in vitro* translation reactions in the presence of 1.5 or 2 μM KAN. The Ec24 strain did not show resistance on agarose plates supplemented with KAN.

In contrast, SQ110 L\TC 23S – A2058G strain kanamycin resistance was provided by SQ110 plasmid, which carries a selection cassette that encodes aminoglycoside 3’-phosphotransferase II enzyme (*35*) that inactivates KAN by phosphorylation in cells. RAPPL-isolated ribosomes from SQ110 L\TC 16S – A1408G strain were the only ribosomes capable of synthesizing eGFP reporter in the presence of 1.5 or 2 μM KAN (Figure 5D). The L\TC 16S – A1408G ribosomes could synthesize eGFP reporter only in the control conditions (no antibiotic present) or in the presence of KAN, confirming the A1408G 16S rRNA mutation responsible for KAN resistance. Interestingly, Ec13 RAPPL-isolated ribosomes did not show any antibiotic resistance in *in vitro* translation assays (Figure 5D). This contrasts sharply with resistance assessed by agarose plates supplemented with antibiotics, where Ec13 cells demonstrated strong resistance to KAN and CLI, and partial resistance to ERY (Figure 5C). We conclude that Ec13 can grow on bacterial agar plates supplemented with KAN, CLI, and ERY, ostensibly due to its secondary multidrug transporter, not ribosome-associated resistance mechanisms (Figure 5C and 5D).

Our results indicate that *E. coli* RAPPL-isolated ribosomes are translationally competent and allow the flexibility of on- or off-bead reaction setup. Therefore, RAPPL can be used to rapidly screen for ribosome or translation factor-associated resistance mechanisms, with plate-based assays confirming alternative mechanisms. Bacterial strains – lab, clinical, or otherwise – that can be cultured or isolated even in relatively small quantities are accessible for study. These data show the ability of RAPPL to isolate and study translation-associated antibiotic mechanisms of resistance from a clinical setting.

### RAPPL generates high-quality materials for structural determination

Finally, we sought to determine the compatibility of RAPPL with structural applications and whether the ribosomes-isolated would be of sufficient quality for structural determination using cryo-electron microscopy. We applied RAPPL to *Cryptococcus neoformans* cells grown in exponential phase isolated in the presence of cycloheximide. *C. neoformans* was selected for two reasons: 1) an 80S structure had yet to be determined, and 2) to determine if the anionic polysaccharides heavily present in the cell wall that are released into the lysate during bead-beading would interfere with RAPPL purification. A small portion of RAPPL eluate was first examined by TEM to see the uniformity of the collected and purified sample (Figure 6A). Since, isolated *C. neoformans* ribosomes represented a majority of the particles in TEM images and had an adequate concentration, the RAPPL eluate was applied to carbon-coated cryo-EM grids, and 2,498 images were collected (Supplementary Figure 10). From an initial 646,683 particles, 399,114 particles were used for 2D classification into 50 classes, and 294,300 particles were refined to ultimately generate a 2.7 Å map with the material eluted from RAPPL straightforwardly (Figure 6 and Supplementary Figure 11).

**Fig. 6:**
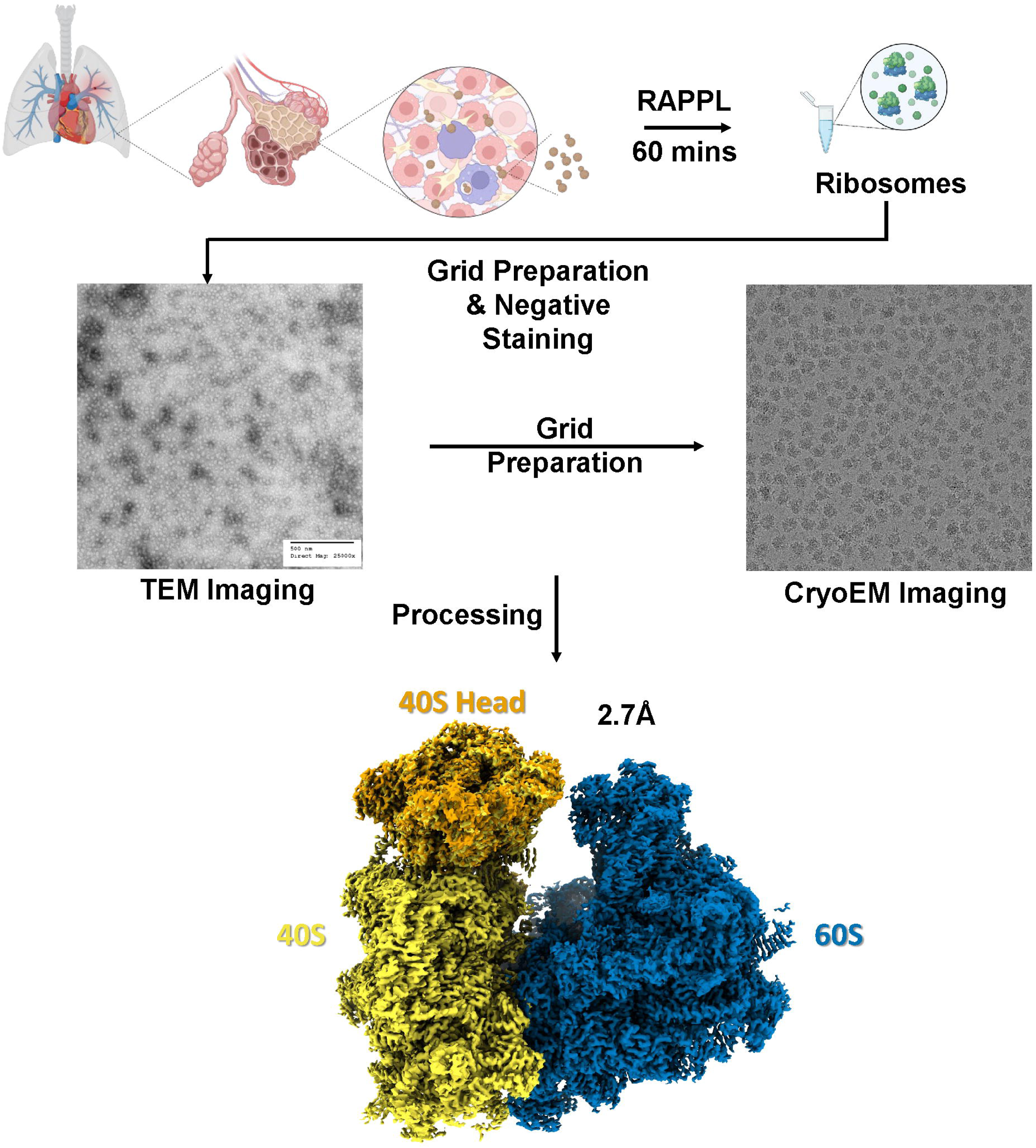
Structural determination of RAPPL products can produce high-resolution CryoEM Maps. *C. neoformans* cells (∼10^8^) in exponential phase were lysed and the clarified lysate used in RAPPL. The ribosomes were eluted in 30 μL of elution buffer. The Eluate was first screened using TEM. Subsequently, grids were prepared using 5 μL of eluate. Movies were captured on FEI Titan Krios G3 300kV Cryo-TEM with Falcon IV Direct Electron Detection camera. Data was processed using cryoSPARC resulting in an ∼2.7 Å global resolution.

This Cryo-EM structure represents the first 80S ribosome structure of *C. neoformans*, an encapsulated yeast causing cryptococcosis and potential death to immuno-compromised and immunosuppressed individuals through infection of the lungs and brain. Moreover, this 2.7 Å structure of 80S ribosomes isolated from 1×10^8^ *C. neoformans* cells fundamentally indicates that the RAPPL method can be used to rapidly isolate ribosomes from biological material, cost-friendly, and efficiently for high-resolution structural studies.

## Discussion

The isolation of ribosomes from different cell types, organs, and organisms imposes several challenges, primarily due to the unique biochemical environment and cellular compositions. The most common problem associated with ribosome isolation is contaminant presence, which can affect RNA quality and protein content during isolation. Different cell/tissue types require different adjustments based on the cell type. Further, the cellular structure of the tissue can affect ribosome isolation; tissues with dense cellular arrangements, such as brain or heart tissue, can be more challenging to homogenize, leading to inefficient lysis and ribosome extraction. Lastly, different cell types/tissues require specific pH levels and ionic conditions for optimal ribosome extraction. Maintaining such conditions during isolation is critical, as deviations can reduce ribosomal activity and purity. Traditional procedures are time-consuming, complex, and often not suitable for all tissue types, especially when rapid isolation is needed for downstream application. Maintaining RNA integrity or achieving high yield and purity simultaneously can be challenging. Methods that obtain purity may reduce yield, making it difficult to obtain ribosomes that are both functional and free of contaminants. Notably, the activity of isolated ribosomes can vary depending on the tissue and isolation method used. In summary, the isolation of ribosomes from different cells/tissues is fraught with challenges that stem from tissue-specific characteristics.

We developed a rapid, facile, and parsimonious method to purify ribosomes and associated factors to overcome these challenges. RAPPL is a robust and versatile method capable of enriching ribosome-associated materials from a wide range of sample types, even those in low abundance and cellular compartments. The downstream application of isolated ribosomes can go in many directions, such as translational studies, drug development, functional analysis, ribosome profiling, structural biology, and posttranslational modification studies. We were able to demonstrate the compatibility of RAPPL with several of these. Our method is well-suited to the currently available technologies and methodologies (Figure 4A & 4B). RAPPL can also be used to study the effects of translational inhibitors on eukaryotic ribosomes (Figure 4B), which could be coupled with factor-specific tandem purifications for further analysis (Figure 4C). Using various *E. coli* strains, including patient-isolated UPEC strains, we can isolate ribosomes via RAPPL and subsequently determine translation-associated mechanisms of antibiotic resistance (Figure 5C and 5D).

The ability to isolate translational machinery from limited starting materials would prove advantageous for those fields in which this is a major limiting factor (e.g., patient samples, clinical isolates, and several parasites). Here, we demonstrate the ability of RAPPL to isolate ribosomes from as little as 5,000 - 10,000 mammalian cells (Figure 2B and Supplementary Figure 5). However, mammalian cells harbor significantly more ribosomes than other organisms. We further demonstrate that our method can isolate and detect ribosomes from as low as one million *P. falciparum* NF54 cells (Figure 2C and Supplementary Figure 7). This is significantly less than necessary for other methods used to study ribosomes and protein synthesis. Often, ∼10^8^ *P. falciparum* cells or more are used in other methodologies like polysome profiling, which depends on gradient volume, with larger gradient sizes requiring up to five times more material. This limitation prevents the use of many clinically relevant organisms and patient biopsies in such studies. While RAPPL will not replace current methods used to study ribosomes and translation, the lower cellular threshold enables researchers to gain access to this data for those organisms and clinical samples that cannot be isolated in sufficient quantities for said traditional methods.

RAPPL purified ribosomes from a wide range of organisms with minor modifications to cell lysis when necessary. This method worked with single celled organisms – intracellular and extracellular – and cultured cells, as well as tissues and whole organisms (Figure 3). Processing single-celled organisms and cultured cells is done similarly. Lysis of these organisms is done using detergent (triton-X100). For those that have cell walls (bacteria, yeast) or multiple membranes (*P. falciparum*), bead-beating is introduced to ensure membrane rupture. In the case of the intracellular parasite *T. gondii*, the host cells are lysed by shearing prior to parasite lysis. *P. falciparum* requires lysis in potassium acetate to ensure ribosome release from the endoplasmic reticulum (*30, 39*), which must then be diluted (1:8) to enable ribosome binding to the beads. This dilution did not prevent enrichment by RAPPL (Figure 3A). However, these lysis methods are known and currently used in their respective fields. Tissues, such as the perfused mouse organs used here, and whole organisms require a breakdown of the tissue structure and cell wall by flash-freezing and subsequent bead-beading or milling to ensure the release of the cytoplasmic contents (Figure 3B). Whole organisms with complex tissue organization, like the *D. rerio* used here, require sample scoring prior to flash-freezing and milling (Figure 3B). Again, these methods are already employed, demonstrating RAPPL’s adaptability for a multitude of sample specimen types (Figure 3). Of import are the conditions under which RAPPL lysis and binding are performed regarding those necessary to maintain ribosome subunit association with mRNA, should this be desired, as well as those needed for other translation-associated factors.

The study of mitochondrial ribosome dysfunction is of clinical importance with a host of life-threatening outcomes (*40*). Using RAPPL, we were able to purify mitochondrial ribosomes rapidly (Figure 3C and Supplementary Figure 8). Our method could be combined with current laboratory or clinical studies to examine mitochondrial ribosomes for functional, composition, and structural analysis (*15, 16*). Although we prioritized using RAPPL for the study of protein synthesis, the method can purify other pertinent ribosome-associated activities, such as ribosome biogenesis (Figure 3C and Supplementary Figure 8). Ribosome biogenesis is essential to the cell cycle (proliferation, differentiation, apoptosis, et cetera), cell and organismal development and plays roles in malignant cell transformation and therapeutic resistance (*41–43*). The study of ribosome biogenesis also provides insights into microbial diversity through ribosome evolution, function, and the development of therapeutic resistance (*9, 44*). The ability to quickly and efficiently harvest this material enables study in these areas, which we were able to demonstrate (Figure 3C and Supplementary Figure 8). Thus, RAPPL enables the purification and study of ribosomes from various cellular compartments, not only cytosolic ribosomes.

Functional and structural analysis of purified ribosomes can provide insight ranging from the effects of drug treatments on the ribosome translation cycles to the outcomes of different cell stressors. Using *in vitro* protein synthesis kits, we were able to visually demonstrate that ribosome organization is maintained by RAPPL (Supplementary Figure 9). We were also able to show that the effects of translation inhibitors on this organization can be visualized using our method (Figure 4A), suggesting that further study of such drug treatments or other stress factors is possible. However, it should be noted, as previously mentioned, that adaptations may be necessary for more nuanced investigation, such as any specific conditions to ensure accessory protein binding and high concentrations of anionic compounds that disrupt poly-lysine: RNA interactions will reduce, if not inhibit, purification by RAPPL.

The compatibility of RAPPL with current technologies like *in vitro* kits and methodologies like polysome gradient profiling provides further flexibility. Enriching from polysome profiling fractions via magnetic bead isolation (Figure 4B) without the necessity of genetic manipulation to introduce affinity tags enables researchers to quickly and freely pursue various avenues of study. It also reduces the time to use isolated products, thereby reducing degradation or complex dissociation that may occur during long centrifugations. Product enrichment using RAPPL over loss often seen with sucrose cushion or gradient centrifugation (*20*) is also a benefit, requiring less starting sample, and RAPPL elution can be used for further purification in tandem with affinity tags associated with the nascent polypeptide chain (Figure 4C), or ribosome or translation associated factors. Purifying and enriching ribosomes is useful for their study in various conditions. However, enriching functioning ribosomes can provide significantly more information through *in vitro* translation studies. Our results indicate that RAPPL products are functional and can be used on (Figure 5A) or off bead (Supplementary Figure 10B), as well as following fluorescence of reporter genes in plate reader assays (Figure 5B) during *in vitro* protein synthesis. This method drastically shortens the time necessary for isolation and functional testing of the isolated ribosomes from bacterial cells, providing a good basis for developing *in vitro* translation kits or lysates from other cells and organisms.

We also demonstrated the clinical applications of RAPPL to study ribosome-associated resistance mechanisms by using clinical UPEC isolates, with mutagenized and lab strains as controls (Figure 5C and 5D). These methods can then be further adapted to plate-based assays (as indicated by Figure 5B)), which lends to the possibility of high throughput assays using ribosomes isolated from various pathogenic organisms on compound libraries. Furthermore, with the right supplementation (i.e., S100 fraction), *in vitro* protein synthesis studies may be possible with eukaryotic organisms.

The purification of ribosomes and ribosomal complexes for structural determination can be quite time and labor-intensive. Here, we demonstrate the ability to obtain cryoEM-ready samples using RAPPL in approximately one hour, producing high-quality structure maps (Figure 6 and Supplementary Figure 11). This process typically requires a significant number of cells, which are sometimes hard to obtain, as is the case with many clinically relevant organisms like the parasites *P. falciparum* or *C. parvum*. To isolate certain ribosomal complexes, such as the pre-initiation complex, more material will be needed, followed by polysome profiling and isolation from the desired sucrose fraction(s) by lengthy ultracentrifugation. Throughout this process, sample loss to handling, degradation, and complex dissociation inevitably occurs. RAPPL provides a means of rapid sample enrichment, which can be performed instead of, prior to, or following polysome profiling, depending on what ribosomal complexes are sought. These options can decrease the required starting sample and/or sample loss at key bottlenecks while reducing the time from lysis to grid preparation and, ultimately, structural determination.

To summarize, the development of methods like polysome profiling has been instrumental in furthering our understanding of ribosomes, protein synthesis, and gene regulation. However, this method has some limitations, such as meeting cell material requirements, lengthy centrifugation times, costly equipment, and sucrose contamination in fractionated samples. To reduce and circumvent these limitations, we present RAPPL, a method defined by its ease of use, wide range of applications, and adaptability. Using RAPPL, we are able to isolate ribosomes as well as ribosome- and translation-associated factors from a wide variety of specimens and in limited ribosome or cell numbers. As shown by mass spectrometry, RAPPL significantly enriches ribosome- and translation-associated factors. Ribosome organization is also maintained during purification and can be visualized by TEM, demonstrating the effects of the addition of mRNA on in vitro protein synthesis kits or mRNA translation inhibitors on mammalian cell lysates. Ribosomes isolated by RAPPL can be used in functional studies, as we did here, showing ribosome-associated resistance mechanisms using *in vitro* protein synthesis kits. This suggests the ability of RAPPL to generate ribosomes for *in vitro* protein synthesis from virtually any organism, given that the right additional factors are supplied, such as those in the S100 fraction. RAPPL is compatible with current methods, enabling ribosome and ribosome-bound protein enrichment from polysome profiling fractions while removing sucrose. RAPPL eluates are also of sufficient quality for structural studies and are capable of producing high-resolution maps by cryoEM for structural determination of ribosomes and ribosome-associated complexes. We hope RAPPL provides researchers with a new means or further flexibility in their ribosome and protein synthesis studies.

## Methods

### Cell Culture and Animal Husbandry

#### Escherichia coli

*E. coli* DH5α cells, uropathogenic patient isolate *E. coli* strains Ec13 and Ec24 (A kind gift from Dr. Jeffrey Henderson (*32*) and *E. coli* rRNA mutagenized lines SQ110 L\TC 16S – A1408G and SQ110 L\TC 23S – A2058G (*35*)(a kind gift from Dr. Nora Vazquez-Laslop and Dr. Alexander Mankin) were cultured overnight in Luria-Bertani (LB) medium. From this overnight culture, 2 mL was used to inoculate 50 mL of LB medium. For mass spectrometry, *E. coli DH5*α cells were grown for 1.5-2 hours. Otherwise, all cells were incubated for 3 hours at 37℃ while shaking at 200 rpm. In the case of rRNA mutagenized lines, cell culture was doubled to 100 mL of LB medium (4 mL inoculation) as these *E. coli* strains grew at approximately half the rate of the other lines.

*E. coli* serial dilution tests were performed on LB agar plates (TEKNOVA LB broth and agar) with varying concentrations of antibiotics until the most effective concentrations of each antibiotic were found—100 mg/mL of erythromycin (Sigma Aldrich) and clindamycin (Sigma Aldrich), 12.5 mg/mL of kanamycin (Gold Biotechnology) and chloramphenicol (Gold Biotechnology). Cultures for each *E. coli* sample were grown overnight without antibiotics in the LB media, and serial dilution tests were performed the next day. Before serial dilutions, overnight cultures’ optical density was equalized to 2.0 within 0.1 at 600 nm (OD_600_) using a Thermo Fisher NanoDrop. After equalizing overnight cultures, serial dilutions of 10x, 100x, 1000x, and 10000x were dropped on plates in 7 µL increments. Plates were then placed in an incubator at 37°C overnight and imaged the following day using a BIO-RAD ChemiDoc imaging system.

### HEK-293, Human Dermal Fibroblasts, and HeLa Cells

HEK T-RExTM-293 cells (R71007, Thermo Fisher), human dermal fibroblasts (HDF, ATCC, PSC-201-012) and HeLa cells (ATCC, CRM-CCL-2) were maintained in DMEM (Gibco) supplemented with 10% heat-inactivated FBS (Gibco), 1×Penicillin Streptomycin and Glutamine (Gibco) and 1 × MEM Non-Essential Amino Acids (Gibco). Cells were grown to 90% confluency and divided 1 to 4 for HEK-293 and HeLa, or 1 to 3 for HDFs for continued growth. CRISPR/Cas9 engineered HEK-293 cell lines (uS-4-Flag, uS-13-Flag, and uL-4-HA tagged HEK-293 cells, Genscript) were treated as HEK T-RExTM-293 cells. Cells were incubated with translation inhibitors 20 minutes before collection when indicated. Cells were collected by trypsinization and washed in 1xDPBS (Thermo Fisher # 14190144; with additional inhibitors when noted) by centrifugation at 500xG for 5 minutes prior to lysis.

#### Plasmodium falciparum

Parasites were cultured as previously described (*45*). Briefly, *P. falciparum* Dd2 or NF54, as well as engineered *Pf*RACK1-mNeonGreen-HA, were maintained by continuous culture at 2-5% hematocrit in human erythrocytes with malaria culture medium (RPMI 1640 supplemented with 5 g/L Albumax II (Gibco), 0.12 mM hypoxanthine (1.2 ml 0.1 M hypoxanthine in 1 M NaOH), and 10 μg/ml gentamicin). Cultures were grown statically under hypoxic conditions in a candle jar atmosphere. Synchronization was done using 5% sorbitol treatment and magnetic purification using MACS cell separation magnets over LD columns. The parasites were washed with 1X PBS prior to lysis.

### Plasmids and genetic modification of *P. falciparum*

The yPM2GT donor vector and Cas9+sgRNA expression plasmid (pAIO3) used to edit *P. falciparum* RACK1 locus has been described previously (*46–48*). For *in-situ* C-terminal tagging of *PfRACK1* with mNeonGreen-3xHA tag, 712 bp immediately upstream of the stop codon (left homologous region [LHR]) and 798 bp of the 3’ UTR (right homologous region [RHR]) was amplified from NF54^attB^ genomic DNA using primers pairs p1-p3 and p4-p5 respectively. The LHR and RHR amplicons were sequentially cloned into the yPM2GT donor vector between AflII/NheI and XhoI/AflII sites, resulting in the plasmid sPL6. Two guide RNA target sites were selected, and the complementary sense and antisense oligos for each sgRNA were annealed and ligated into the AflII site of the pAIO3 plasmid using an In-Fusion cloning kit (Takara). Before transfection, the donor plasmid sPL6 was linearized using AflII and co-transfected into *P. falciparum* NF54^attB^ cells with the Cas9+sgRNA plasmids designed to target *PfRACK1*. Transgenic cells were selected with 2 μM DSM1, and the expected integration was confirmed by diagnostic PCR using p10-p11, p12-p13, and p10-p13 primer pairs.

#### Toxoplasma gondii

*T. gondii* ME49 parasites were continuously cultured in human foreskin fibroblast (HFF) cell monolayers as previously described (*49*). HFF cells were maintained in Dulbecco’s modified Eagle’s medium (Invitrogen) supplemented with 10% HyClone fetal bovine serum (GE Healthcare Life Sciences), 10 μg/mL gentamicin (Thermo Fisher Scientific) and 10 mM glutamine (ThermoFisher Scientific) (D10). *T. gondii* parasites were isolated from host cells as previously (*50*). Briefly, parasites were cultured to high parasitemia (∼75%) in two T25 flasks. The monolayers were scraped and combined in 10 mL of D10 medium. The cell suspension was passed through 22G blunt-end syringe 3 times to disrupt host cells. Host cell debris was filtered out by passing through a pre-wet 3 μm polycarbonate membrane, which was then washed with and additional 5 mL of D10 medium. The freed T. gondii cells were pelleted by centrifugation (400 x g for 10 mins). The parasites were washed with 1X PBS prior to lysis.

#### Cryptosporidium parvum

Purified *C. parvum* oocysts were graciously provided by the Sibley Lab per the lab protocol (*51*). Oocysts (10^7^) were bleached by treating them with 40% bleach and incubating them on ice for 10 mins. The oocysts were removed by centrifugation (900 x g for 3 mins at 4). The supernatant was removed, and the oocysts were washed three times with 1X DPBS + 1% BSA. Excystation was performed by combining equal volumes of resuspended oocysts (100 μL) and 1X DPBS + 1.5% sodium taurocholate. The oocysts were then incubated for 60-75 minutes at 37℃. Excystation was confirmed by brightfield microscopy (∼80%). The parasites were centrifuged for 3 mins at 1400 x g and washed with 1X DPBS twice prior to lysis.

#### Saccharomyces cerevisiae

An overnight culture of S. cerevisiae was grown by inoculating 10 mL of yeast-peptone-dextrose (YPD) growth medium with 200 μL of glycerol stock at 30. The culture was harvested by centrifugation at 3500 x g for 5 mins at 4. The culture was washed with 1X PBS prior to lysis.

#### Cryptococcus neoformans

*C. neoformans KN99*α were grown and generously provided by Dr. Tamara Doering’s lab (Washington University in St. Louis). Briefly, cultures were grown on yeast extract-peptone-dextrose (YPD) plates for two days at 30. YPD liquid medium was inoculated with single colonies and grown overnight at 30 while shaking at 230 RPM. Overnight cultures were diluted to an OD600 of 0.2 and grown to 0.6 (exponential phase). Cultures were pelleted and washed with 1X PBS prior to lysis.

#### Caenorhabditis elegans

*C. elegans* worms were generously provided by Dr. Tim Schedl’s lab (Washington University School of Medicine). A synchronized population of 2000 - 3000 *C. elegans* embryos were obtained following a two-hour egg lay on NGM plates seeded with *E. coli*. Worms were grown continuously to the young adult stage and harvested before fertilized eggs appeared. Worms were washed 4-times in M9 buffer and then 2-times in deionized, distilled water, pelleted and frozen in liquid nitrogen prior to lysis.

#### Danio rerio

Zebrafish (D. rerio wild-type AB) were graciously provided by the Stratman lab, and experimental procedures were done per approved guidelines by the Washington University in St. Louis School of Medicine Institutional Animal Care and Use Committee (IACUC). Fish were euthanized by an ice water bath (5 parts ice/1 part water, 0-4) separated from ice chips by a fine mesh strainer for a minimum of 10 minutes after cessation of opercular movement. Animals were snap-frozen in liquid nitrogen and stored at -80°C until further use.

#### Mus musculus

Perfused mouse organ sections from WT C57Bl6/N mouse (Charles River, Cat#: C57BL/6NCrl were generously provided by the Dr. Maxim Artyomov lab (Washington University School of Medicine) and experimental procedures done per approved guidelines by the Washington University in St. Louis School of Medicine Institutional Animal Care and Use Committee (IACUC). Mice were humanely euthanized in a CO_2_ chamber. Individual organs (liver and spleen) were removed and snap-frozen in liquid nitrogen. Frozen tissue has been stored at -80°C until further use.

### RNA Poly-lysine Affinity Purification (RAPPL)

Cells, organs, and organisms were maintained as mentioned above. Cells were centrifuged, their growth mediums removed, washed with PBS, and transferred to 2.0 mL microcentrifuge tubes. Cells were then resuspended in RAPPL lysis buffer (100 mM HEPES·KOH solution, pH 7.5, 50 mM KCl, 10 mM Mg(OAc)_2_, 1% Triton-X, 1 mM DTT, Protease Inhibitors (Cell Signaling), 40 U/mL RNaseOUT, 20 U/mL ^Superas·IN™^ RNase Inhibitor, 4 U/mL DNase I). Bacterial and fungal samples were resuspended in a ratio of 1:2:1 cell pellet: lysis buffer:acid-washed glass beads (Sigma #G8772). Mammalian cell lines and *T. gondii* were resuspended in 500 μL to 1ml of lysis buffer. *P. falciparum* and *C. parvum* were resuspended in 400 μL and 250 μL modified lysis buffer (25 mM K-HEPES, pH 7.5, 400 mM KOAc, 15 mM Mg(OAc)_2_, 2% Triton-X100, 1 mM DTT, Protease Inhibitors (Cell Signaling), 40 U/mL RNaseOUT, 20 U/mL Superas·IN™ RNase Inhibitor, 4 U/mL DNase I). Following lysis, samples in modified lysis buffer were diluted 8X in RAPPL binding buffer (100 mM HEPES-KOH pH 7.5, 10 mM Mg(OAc)_2_, 1 mM DTT, Protease Inhibitors (Cell Signaling), 40 U/mL RNaseOUT, 20 U/mL Superas·IN™ RNase Inhibitor, 4 U/mL DNase I). Perfused organ sections were homogenized and resuspended in 1 mL RAPPL lysis buffer, sonicated two times for 15s at 60 Hz before further bead-beating. For all single-celled organisms, *C. elegans*, and perfused mouse organ sections, bead-beating was performed using a BeadBug™ 3 Microtube homogenizer (Benchmark Scientific) for 30s at 4,000 Hz, three times at 4. For mammalian cell lines, no bead-beating was performed. Following euthanasia, the *D. rerio* sample was finely scored and segmented using a scalpel and then flashed frozen by plunging into a liquid nitrogen bath. The frozen sample was then pulverized into a fine powder using a mixer miller MM 400 (Retsch). This powder was resuspended in 1 mL RAPPL lysis buffer and incubated for 15 mins, rotating end-over-end at 4. The lysates were clarified by centrifugation for 10 mins, 21,100 x g, 4. Lysates were transferred to a new tube and centrifugated again for 5 mins, 21,100 x g, 4. Lysates were then applied to 100 μL polylysine magnetic beads (Molecular Cloning Laboratories) for 15-30 minutes at 4, rotating. Beads were removed from the flow-through by a magnet. The beads were washed three times with 250 μL RAPPL wash buffer (100 mM HEPES·KOH solution, pH 7.5, 50 mM KCl, 10 mM Mg(OAc)_2_, 1 mM DTT, Protease Inhibitors (Cell Signaling), 40 U/mL RNaseOUT, 20 U/mL Superas·IN™ RNase Inhibitor). Beads were then eluted with 50-200ul RAPPL elution buffer (100 mM HEPES·KOH solution, pH 7.5, 50 mM KCl, 10 mM Mg(OAc)_2_, 2 mg/mL poly-D-glutamic acid (Sigma #4033), 1 mM DTT, Protease Inhibitors (Cell Signaling), 40 U/mL RNaseOUT, 20 U/mL Superas·IN™ RNase Inhibitor) incubating for 15 mins at room temperature or 4 with rotation or agitation to maintain bead suspension.

### Tandem Purifications

RAPPL eluates from lysates of non-induced and arabinose-induced *E. coli Bl21 (DE3)* cells expressing 2xHA-TevC-eGFP were applied to 25µl of αHA magnetic beads for 2 hours at 4, rotating. When noted, lysates and buffers included 50 µM chloramphenicol (CHL). Beads were washed three times using RAPPL wash buffer. Beads were eluted in 50 μL RAPPL wash buffer plus 1µl of TEV protease (TEV Protease His, Genscript) overnight at 4 rotating.

### Cellular Fractionation - Cytoplasmic, mitochondrial, nuclear

Nuclear and cytoplasmic fractions of Flp-In™ T-REx™ 293 with engineered HA-tag in uL-4 were separated according to the manual for NE-PER™ Nuclear and Cytoplasmic Extraction Reagent (Thermo Scientific™). Ribosome enrichment in both cellular and nuclear fractions was analyzed using western blot analysis and an antibody for the HA-tag, as indicated above in the RAPPL section. Nuclear fraction separation was confirmed using western blot analysis and probing with antibody for topoisomerase II β (Thermo Fisher Scientific # A300-950A). Cytoplasmic and mitochondrial fraction separation was done by rapid enrichment of mitochondria by a previously published procedure (*52*). The mitochondrial enriched fraction was lysed using RAPPL lysis buffer, and a standard RAPPL procedure was followed up for ribosome enrichment. Western blot analyses confirmed the successful separation of cytoplasmic and mitochondrial fractions and the enrichment of ribosomes in both fractions. Enrichment of cytoplasmic ribosomes was analyzed using Flag-antibody for endogenously tagged uS-4-Flag 40S ribosomal protein. In contrast, the mRPS35 antibody for mitochondrial ribosomal protein S35 (Proteintech® Cat No. 16457-1-AP) was used for the enrichment of mitochondrial ribosomes and separation of the mitochondrial fraction. RAPPL eluates from cytoplasmic, nuclear, and mitochondrial fractions were further analyzed by TEM, as were other RAPPL isolated samples.

### RNA Quality Analysis

#### Agarose gel

Standard 2% agarose (w/V) gel electrophoresis using Tris-Acetate Buffer with the addition of 0.5% Clorox bleach (*53*) was performed for the analysis of RAPPL isolated RNA species from human cell cultures (HEK293, HeLa, and HDFs). 1kb and 100 base pair DNA ladder Quick Load® markers (NEB # N0468L and N0467L) and 6x Gel loading Dye (NEB #B7024S) were used for sample preparation and as controls. HeLa cell lysate from the 1-Step Human IVT Kit (Thermo Fisher Scientific #88882) and yeast tRNA (Thermo Fisher Scientific #AM7119) were used as controls for rRNA and tRNA species.

### RNA Quality and Quantity Assessment

Total RNA was isolated from *E. coli* cultures using the RAPPL protocol. The RNA concentration and purity of the RNA were initially measured using a NanoDrop spectrophotometer (Thermo Fisher Scientific), and RNA integrity was further assessed using an Agilent 2100 Bioanalyzer (Agilent Technologies). For Bioanalyzer analysis, 1 µL of RNA (approximately 50 ng/µL) was used per sample and mixed with 1 µL High Sensitivity RNA Screen Tape Sample Buffer 96.00 (Agilent, cat. # 5067-5570). ScreenTape Ladder (Agilent, cat. # 5067-5081) was thawed on ice, mixed gently by flicking or vortexing at low speed, and briefly centrifuged to collect contents. The ladder and the sample are denatured at 72C for 3 min and placed on ice for 2 min. A new High Sensitivity RNA ScreenTape (Agilent, cat. # 5067-5579) was inserted into the TapeStation system (Agilent technologies 4200 TapeStation, 2200 TapeStation Controller Software), and 2 µL of each RNA sample was loaded into the corresponding wells of the sample plate. The TapeStation software was then used to initiate the High Sensitivity RNA analysis protocol. Samples were automatically processed, and the software generated quality, quantity, and sizing data.

### Polysome Profiling

For polysome profiling, equal numbers of Hek T-RExTM-293 cells (R71007, Thermo Fisher) were plated and 24 hours later were treated with 100 μg/ml cycloheximide for 15 min before harvesting. A total of 6 × 10^6^ cells were lysed in 500 μL of polysome lysis buffer (10 mM HEPES pH 7.4, 100 mM KCl, 5 mM MgCl2, 1 mM DTT, 1% NP-40, 100 μg/mL cycloheximide, 1X protease inhibitor cocktail, 25 U/mL DNase I, and 20 U/mL RNase Inhibitor) on ice for 15 min before clearing at 13,000 rpm for 10 min at 4 °C. 1.5 mg of the lysate was layered over a 5–50% sucrose gradient (20 mM HEPES, 200 mM KCl, 10mM MgCl2, 1 mM DTT, 100 μg/mL cycloheximide) (BioComp, gradient master 108) and subjected to centrifugation at 35,000 rpm for 2.5 hr at 4 °C using a SW41Ti rotor (Beckman). The polysome profile in sucrose gradients was resolved using a Brandel gradient fractionator. Absorbance was followed at 254 nm (Brandel UA-6). Fractions were pooled and subjected to RAPPL.

### *In Vitro* Translation Assays

#### End-point

All *E. coli in vitro* translation experiments with RAPPL purified ribosomes were performed using New England Biolabs (NEB) PURExpress D Ribosome Kit (NEB #E3313S) and New England Biolabs PURExpress In Vitro Protein Synthesis Kit (NEB #E6800). All *E. coli in vitro* translation experiments used a DNA template created using PCR to amplify the eGFP gene-containing region of a recombinant plasmid containing the eGFP target protein in a pBAD vector. As in accordance with the NEB PURExpress protocol (https://www.neb.com/en-us/-/media/nebus/files/manuals/manuale6800_e3313_e6840_e6850.pdf?rev=ba7a388352b a4d0fb8089268e1852843&hash=9576CB18CA6990DD5925500898ACFB69), the DNA template contained the in-frame coding sequence for the target protein along with a starting codon, stop codon, T7 promoter sequence upstream of the target protein, ribosome binding site upstream from translation region, a spacer region 6 base pairs downstream from the stop codon, and a T7 terminator sequence downstream of the stop codon. A pBAD specific T7 forward primer and pBAD specific polyA tail reverse primer were used for all PCR amplification reactions (T7 forward primer used for amplification was 5’ TAATACGACTCACTATAGGGAGAAATAATTTTGTTTAACTTTAAGAAGGAG 3’, and the pBAD specific reverse primer used for amplification was 5’ TTTTTTTTTTTTTTTTTTTTTTTTTTTTTTTTTTTTTTTTTAAACTCAATGGTGATGGTG 3’). All PCR reactions to create the DNA template were performed using NEB’s Phusion-HF polymerase kit (M0530S). All PCR products were analyzed on 1% agarose gels and purified using Zymo Research’s Zymoclean Gel DNA Recovery Kit (catalog # 11-301).

All reactions were assembled on ice and in accordance with NEB’s protocol for the kit. All recommended concentrations of solutions and reagents was followed for all experiments unless otherwise stated, so all reactions (except for on-bead translation experiments) were incubated at 37°C for 4 hours. On-bead translation experiments were incubated in a table-top Thermomixer at 37°C and 850 RPM for 4 hours. Except for the experiment testing ideal *in vitro* ribosome concentrations, 4.5 mL of RAPPL purified ribosomes with concentrations averaging ∼1.5 mg per mL were used for all other experiments. Concentrations of RAPPL-purified ribosomes used for functional assays were determined by measuring absorbance at 260 nm using spectrophotometry (NanoDrop, Thermo Fisher Scientific). DNA template concentrations were consistent at 150ng of eGFP template per reaction for all end-point experiments. Reactions with antibiotics present had antibiotics added immediately before 37°C incubation.

After the 4-hour incubation, 2x sample buffer with 5% 2-Mercaptoethanol (Sigma Aldrich) was added to each reaction, and samples were boiled at 95°C for 5 minutes before being loaded onto SDS PAGE gels for western blot analysis. Equal amounts of samples were run on SDS PAGE gels (BIO-RAD 4-12% gradient Bis-Tris or XT Precast gels with catalog numbers mples were run on SDS PAGE gels (BIO-RAD 4-12% gradient Bis-Tris or XT Precast gels with catalog numbers #3450123 or #3450124) using XT MES running buffer (BIO-RAD). A semi-dry transfer of the SDS-PAGE gel was carried out using BIO-RAD’s Immun-Blot PVDF membrane before the membrane with transferred protein was blocked in reconstituted (with PBS) 5% nonfat dry milk for 1 hour (Research Products International). After blocking, the membrane with protein was incubated with 1:3000 diluted primary antibody overnight, washed with PBST (PBS plus 0.1% Tween), and then incubated in 1:10000 diluted secondary antibody for 1 hour before imaging using chemiluminescence on a BIO-RAD ChemiDoc imaging system. Antibodies used for experiments include eGFP (Living Colors A.v. Monoclonal Antibody JL-8; catalog # 632381 and anti-mouse HRP-linked secondary antibody (Cell Signaling; catalog #7076).

#### Kinetic Plate Assay

The DNA template for the kinetic plate *in vitro* assays of the RAPPL purified BL21(DE3) (Intact Genomics; catalog# 1051-24) *E. coli* ribosomes was created using the same process as the end-point assay DNA templates, and reactions were assembled on ice and in the same manner and concentrations as the end-point in vitro assays. A BioTek Cytation, 5 imaging reader, was used for all kinetic assays, and eGFP fluorescence was measured in 1-minute intervals for 2.5 hours in a 384 well plate at 37°C using relative fluorescence units with excitation settings at 488 +/- 9 and emission settings at 507 +/- 9 with gain settings set to 122. The BL21 RAPPL purified ribosome replicates each had 200 ng of eGFP tagged DNA template, and the negative control replicates had 0 ng of eGFP tagged DNA template--all other reaction conditions were kept the same between samples.

### Mass Spectrometry

The beads isolated after immunoprecipitation were incubated in 80 µL of buffer (2M urea, 50 mM tris (pH 7.5), 1 mM dithiothreitol, and 5 µg/mL trypsin (Promega: V511C)) for 1 hour at 25C° and 1000 rpm to partially digest the proteins of the bead–generating an initial eluate. Two additional washes (2M urea, 50 mM tris (pH 7.5)) were performed to maximize yield. The initial eluate and washes were combined and clarified by spinning at 5000g.

Following elution, half of the IP eluate was further reduced with 5mM DTT for 30 min at 25C° and 1000 rpm, and alkylated in the dark with 10mM iodoacetamide for 45 min at 25C° and 1000 rpm. For the flow-through (FT) and input samples (IN), 50ug of protein was reduced and alkylated under the same conditions. Samples were diluted with 50 mM tris for a final urea concentration of < 2M. EDTA was added for a final concentration of 10 mM, followed by SDS to 1%.

Magnetic SP3 beads were made by combining equal volumes of carboxylate-modified hydrophilic (Cytiva: 45152105050250) and hydrophobic beads (Cytiva: 65152105050250). Each sample was used to resuspend 500 µg of SP3 beads. 100% ethanol was added to the sample at a 1:1 volumetric ratio to precipitate the protein material onto the beads. The samples were then incubated for 15 minutes at room temperature.

Following incubation, the beads were washed thrice with 1 mL of 80% ethanol and reconstituted in 100 µL of freshly prepared ammonium bicarbonate with 0.5 µg of trypsin. The samples were incubated overnight at 37°C and 700 rpm to digest the proteins of the SP3 beads. Tryptic peptides were dried in a vacuum concentrator and resuspended in 3% acetonitrile/0.2% formic acid for a final 0.25 µg/µL peptide concentration.

#### LC-MS/MS analysis on a Q-Exactive HF

Approximately 1 μg of total peptides were analyzed on a Waters M-Class UPLC using a 15 cm x 75 µm IonOpticks C18 1.7 µm column coupled to a benchtop Thermo Fisher Scientific Orbitrap Q Exactive HF mass spectrometer. Peptides were separated at a 400 nL/min flow rate with a 90-minute gradient, including sample loading and column equilibration times. Data were acquired in data-dependent mode using Xcalibur software; each cycle’s 12 most intense peaks were selected for MS2 analysis. MS1 spectra were measured with a resolution of 120,000, an AGC target of 3e6, and a scan range from 300 to 1800 m/z. MS2 spectra were measured with a resolution of 15,000, an AGC target of 1e5, a scan range from 200–2000 m/z, and an isolation window width of 1.6 m/z.

Raw data were searched against the Homo sapiens and Escherichia coli proteomes (UP000005640 and UP000000625, respectively) with MaxQuant (v2.6.3.0). The ppm of a protein’s iBAQ value was calculated to determine protein enrichment within a sample. This was done by dividing a protein’s intensity by the sum of all protein intensities in the respective sample and multiplying the resulting fractional value by 1,000,000. After that, a pseudocount of +1 was applied before the ppm values were log2-transformed. The log2 values were used to assign each protein a rank within its sample, serving as a. The iBAQ ppm, log2 values, and rank were subsequently used as indicators for protein enrichment within a sample.

### Transmission Electron Microscopy

For analyses of ribosome preparations, samples were allowed to absorb onto freshly glow discharged formvar/carbon-coated copper grids (200 mesh, Ted Pella Inc., Redding, CA)) for 10 min. Grids were then washed two times in dH2O and stained with 1% aqueous uranyl acetate (Ted Pella Inc.) for 1 min. Excess liquid was gently wicked off, and grids were allowed to air dry. Samples were viewed on a JEOL 1200EX transmission electron microscope (JEOL USA, Peabody, MA) with an AMT 8-megapixel digital camera (Advanced Microscopy Techniques, Woburn, MA).

### Cryo-EM

Grid preparation: RAPPL samples were applied to holey carbon, carbon-coated (2 nm thickness) Quantifoil R2/2 300 mesh grids that had been glow-discharged for 15s using an EMS GloQube Glow Discharger, which were then blotted for 2.5s at 4 in 100% humidity. Samples were then vitrified by plunging into liquid ethane and cooled with liquid nitrogen using the Mark IV Vitrobot (FEI, Hillsboro, Oregon). Vitrified samples on grids were stored in liquid nitrogen prior to imaging. Data were collected on a Titan Krios G3 300 kV electron microscope (Thermo Fisher Scientific) with a sample auto-loading system, Cs Aberation Corrector, Volta Phase Contrast System, and STEM detector operating at 300 kV using a with Falcon IV Direct Electron Detection camera. Images were collected with the automated data collection software EPU 3 and processed on-the-fly with cryoSPARC Live (Thermo Fisher Scientific). A total of 2,498 videos with a total dose of 55 e/Å^2^ split over 50 fractions (individual dose: 1.1 e/Å^2^ per fraction) at a nominal magnification of 59,000x with a calibrated pixel size of 1.122 Å. Data were collected with a defocus range of –0.6 to –2.0 μm.

### Cryo-EM Image Processing and Reconstruction

All data processing was done using cryoSPARC (*54*). The collected framesets were corrected for beam-induced motion on-the-fly using cryoSPARC Live patch motion correction and averages of all 50 frames were used for image processing. The contrast transfer function parameters were determined using Patch CTF job. A total of 561,289 particles were automatically picked using the blob-based picker. Particles were extracted and sorted into 100 2D classes, and artefactual particles were removed. Curated particles were then used in template-based particle selection, yielding 646,683 particles, which were again sorted into 50 2D classes. The final curated 2D classes yielded 399,114 particles, which were used for Ab Initio 3D reconstruction with 3 classes. Non-artefactual classes were subject to non-uniform refinement. Masks for the 60S, 40S, and 40S heads were generated using ChimeraX v1.8 (*55*). These were used for focused refinements and particle subtraction. The resolutions reported were based on gold-standard Fourier shell correlation curves.

## Acknowledgments

We are thankful to all the members of Pavlovic Djuranovic, Djuranovic, Hashem, and Jovanovic labs for their help in our study and critical reading of the manuscript. We are thankful to Dr. Nora Vazquez-Laslop, Dr Alexander Mankin, Dr. Hani Zaher, Dr. Tim Schedl, Dr. Yury Polikanov, and Dr. Juan Alfonzo for their valuable comments and careful reading of the manuscript. NIGMS R01GM136823 and R01GM112824, Chen-Zuckerberg Neurodegeneration Initiative, and Siteman Cancer Center Investment Award sponsored the work in Djuranovic Lab. Work in Pavlovic-Djuranovic is supported by NIGMS R01GM136823. The work in Hashem Lab is supported by the European Research Council Consolidator Grant (SPICTRANS ID: 101088541).

We thank Genscript for re-creating CRISPR/Cas9-engineered HEK-293 cell lines. Dr. Sergej Djuranovic, Dr. Slavica Pavlovic Djuranovic, and Dr. Jessey Erath hold a provisional patent on “Method of use, procedures, and application of purified ribosomes and translation material using poly-lysine and other poly-basic polymers.”

