## Supplementary Figures for "A rapid, facile, and economical method for the isolation of ribosomes and translational machinery for structural and functional studies"

#### Supplementary Figure 1

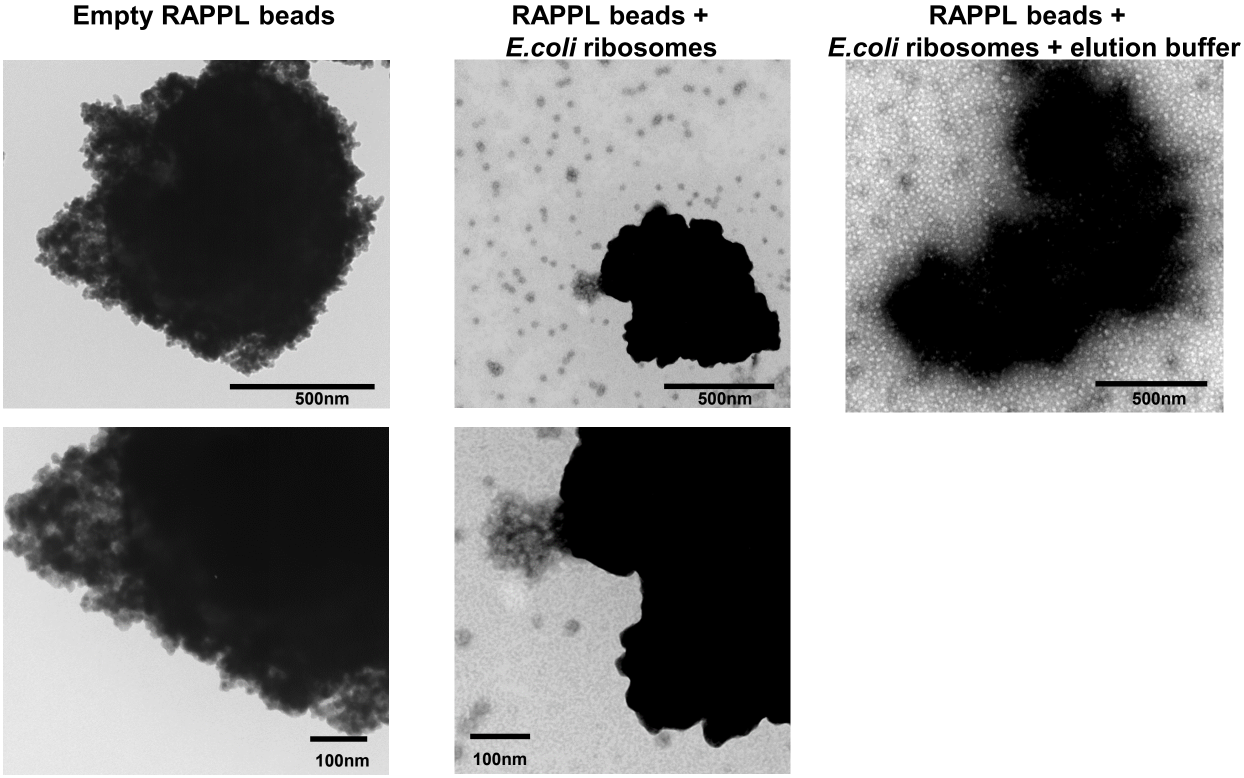


**Supplementary Figure 1. Binding of E.coli ribosomes (NEB PURExpress® *in vitro* kit) to poly-L-lysine beads (Puramag™, MCLAB).** Poly-L-lysine magnetic beads were incubated with wash buffer only or with *E. coli* ribosomes in absence or presence of elution buffer.

### Supplementary Figure 2

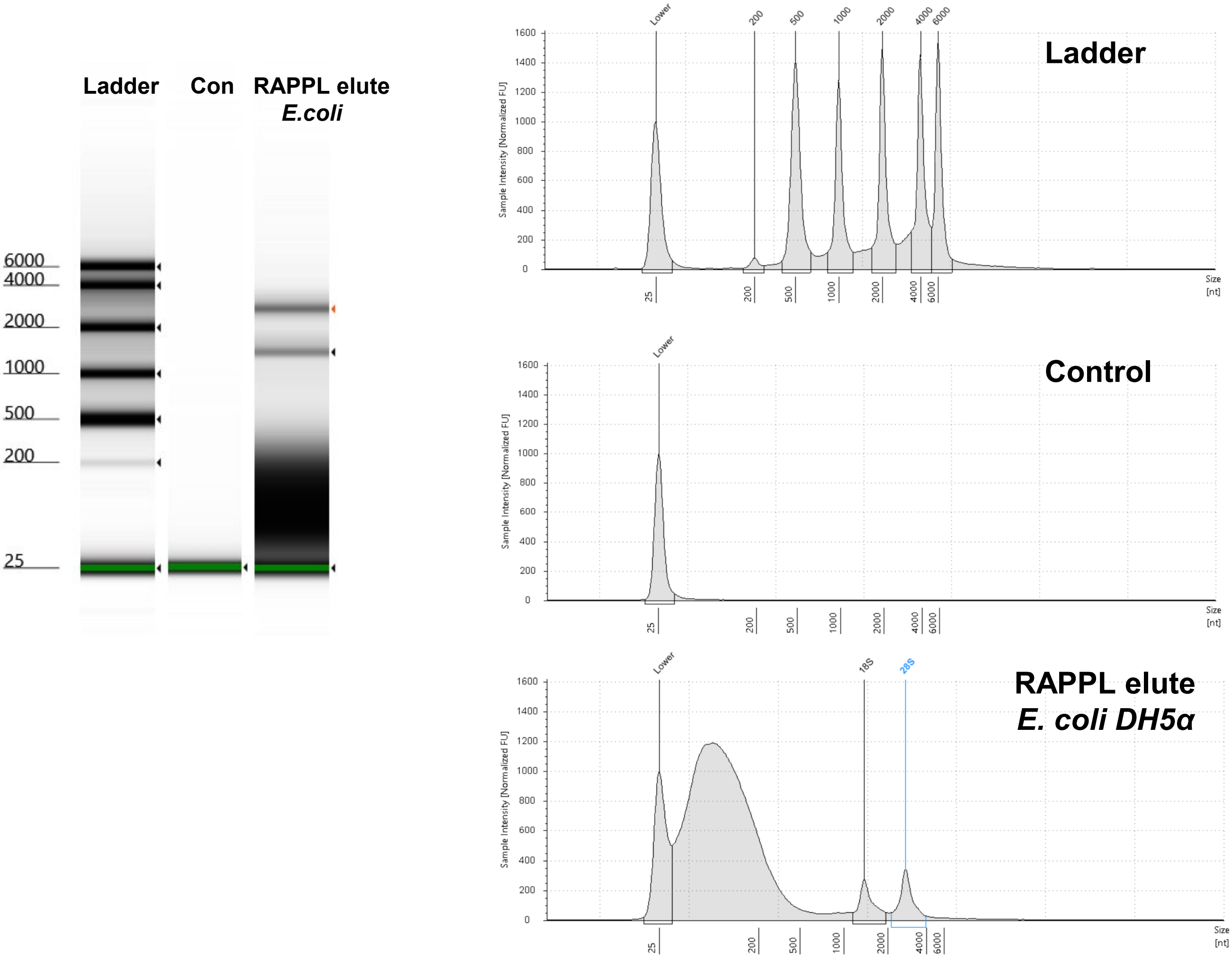


**Supplementary Figure 2. Bioanalyzer analysis of RNA species associated with RAPPL purification from *E.coli* DH5α cells.** Image of the ladder, control (elution buffer) and diluted RAPPL elution from *E. coli* lysate purification. Electropherogram indicating ribosomal subunits as well as smaller and partially degraded RNA species.

### Supplementary Figure 3

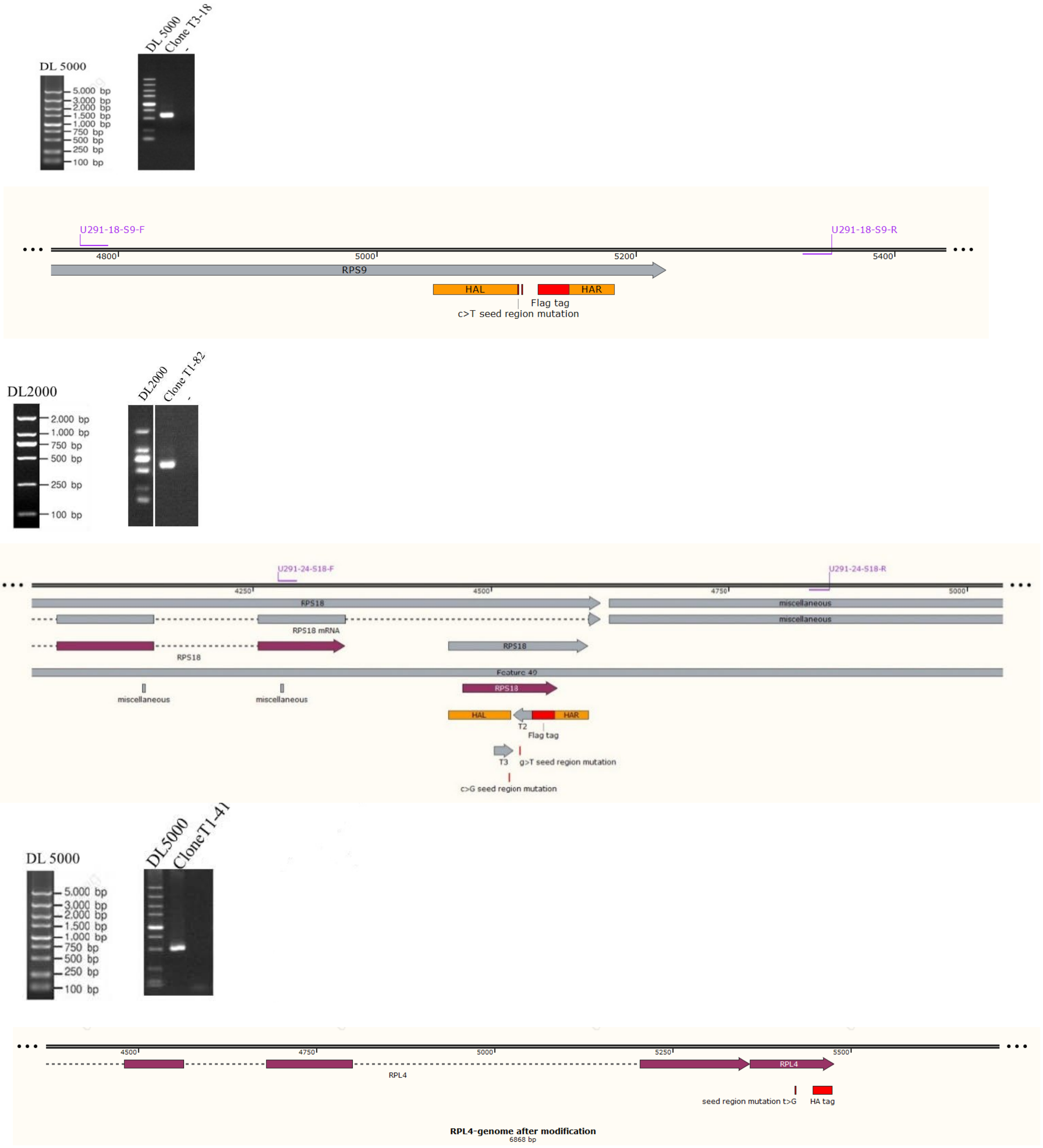


**Supplementary Figure 2. CRISPR/Cas9 engineered Hek293 cell lines.** Images of the DNA isolated from engineered uS4, uS13 and uL4 as well as map of primers for modified single clone validation are indicated. Targeted insertion of Flag- and HA-tags in ribosomal proteins is indicated in maps as well. Cell lines and images were generated by Genscript.

#### Supplementary Figure 4

**A.**

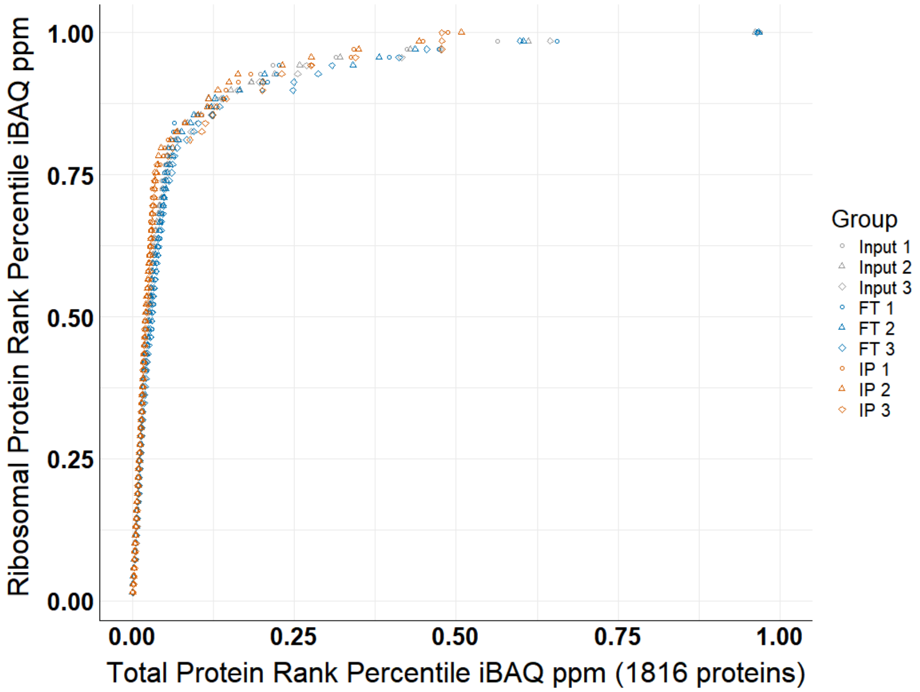


**B.**

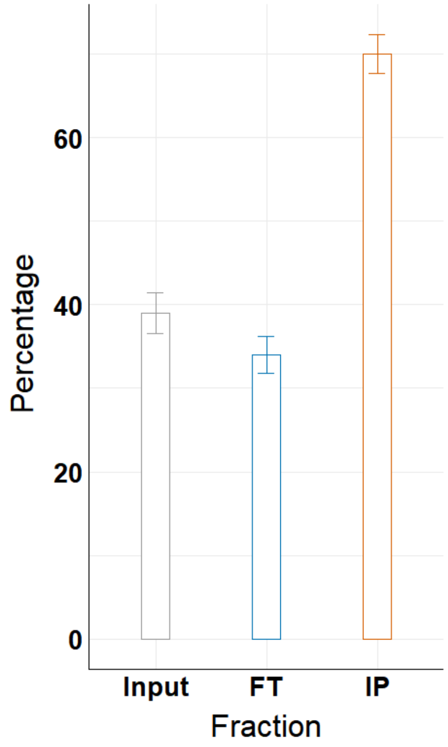


**Supplementary Figure 4. Mass spectrometry analysis of *E. coli* RAPPL purifications. a,** Plot of the of each ribosomal protein's rank percentile in relation to total protein rank percentile for each replicate of input, flow-through (FT), and bead-bound (IP) fractions. **b,** Graph representing percentage of ribosomal proteins in total protein associated with input, flow-through (FT), and bead-bound (IP) fractions. Error bars represent standard deviation of triplicate averages.

### Supplementary Figure 5

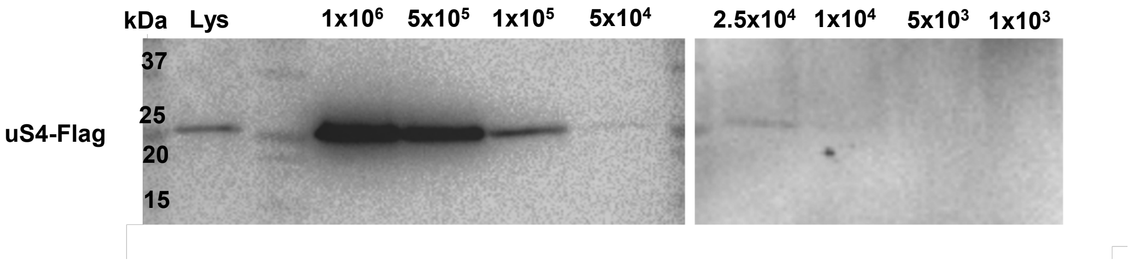


**Supplementary Figure 5. RAPPL can purify ribosomes from limited HEK-293 cell sample.** Western blot analysis using αFlag antibody on RAPPL eluates of HEK-293 cells in which uS4 was Flag-tagged by CRISPR/Cas9 whereby the starting cells were diluted to $1x10^6$, $5x10^5$, $1x10^5$, $2.5x10^4$, $1x10^4$, $5x10^4$ and $1x10^3$ cells prior to lysis. Molecular weight markers indicate size of tagged uS4 protein.

### Supplementary Figure 6

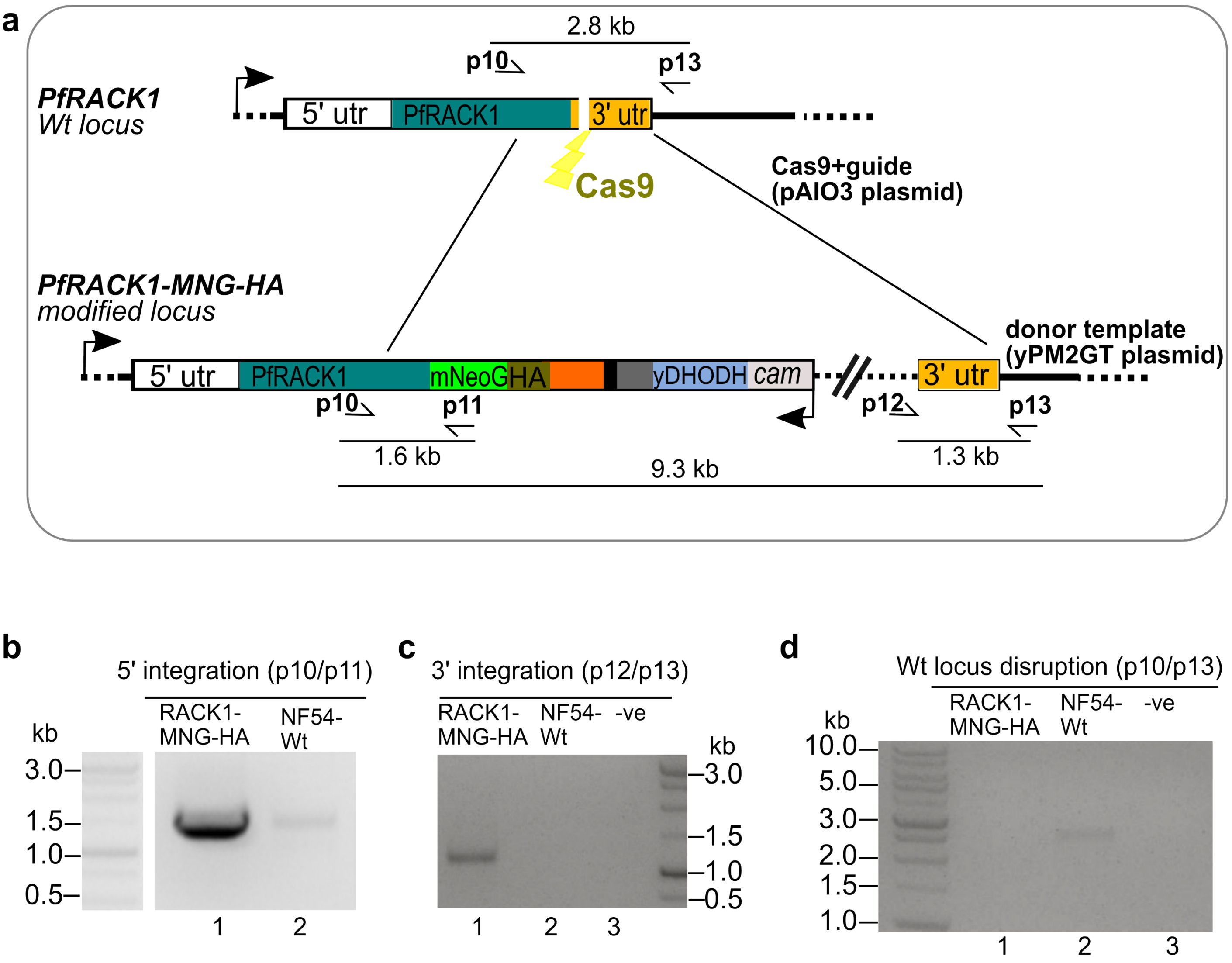


**Supplementary Figure 6:** Endogenous C-terminal tagging of *Plasmodium* PfRACK1 with mNeonGreen-3xHA tag. **(a)** Schematic of CRISPR/Cas9 strategy used to edit *PfRACK1* locus. **(b-d)** Agarose gels showing PCR products from genomic DNA of the cell line with the modified locus, demonstrating PCR amplicons across the 5’ **(b)** and 3’ **(c)** integration junction as well as the absence of the original locus **(d)**. Primers used are as indicated in **(a)** and are listed in material section. MNG: mNeonGreen; HA: hemagglutinin epitope tag; utr: untranslated region; yDHODH: yeast dihydroorotate dehydrogenase selectable marker; cam: calmodulin promoter; NF54-Wt: wild type parental line; -ve: no template negative control.

#### Supplementary Figure 7

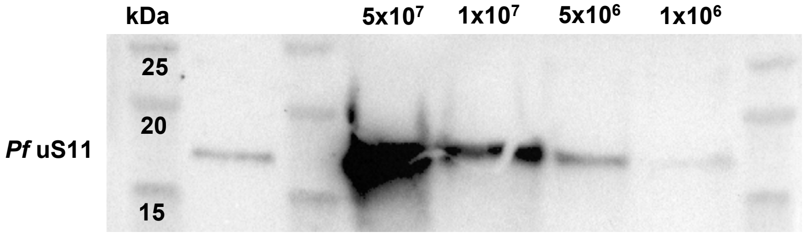


**Supplementary Figure 7. RAPPL can purify ribosomes from limited P. falciparum.** Western blot analysis using α*Pf* uS11 antibody on RAPPL eluates of P. falciparum NF54 cells. Starting cells were diluted to $5x10^7$, $1x10^7$, $5x10^6$, and $1x10^6$ cells before RAPPL procedure was executed. 20% of total bound fraction was used for analysis. Molecular weight markers indicate size of tagged Pf uS11 protein.

#### Supplementary Figure 8

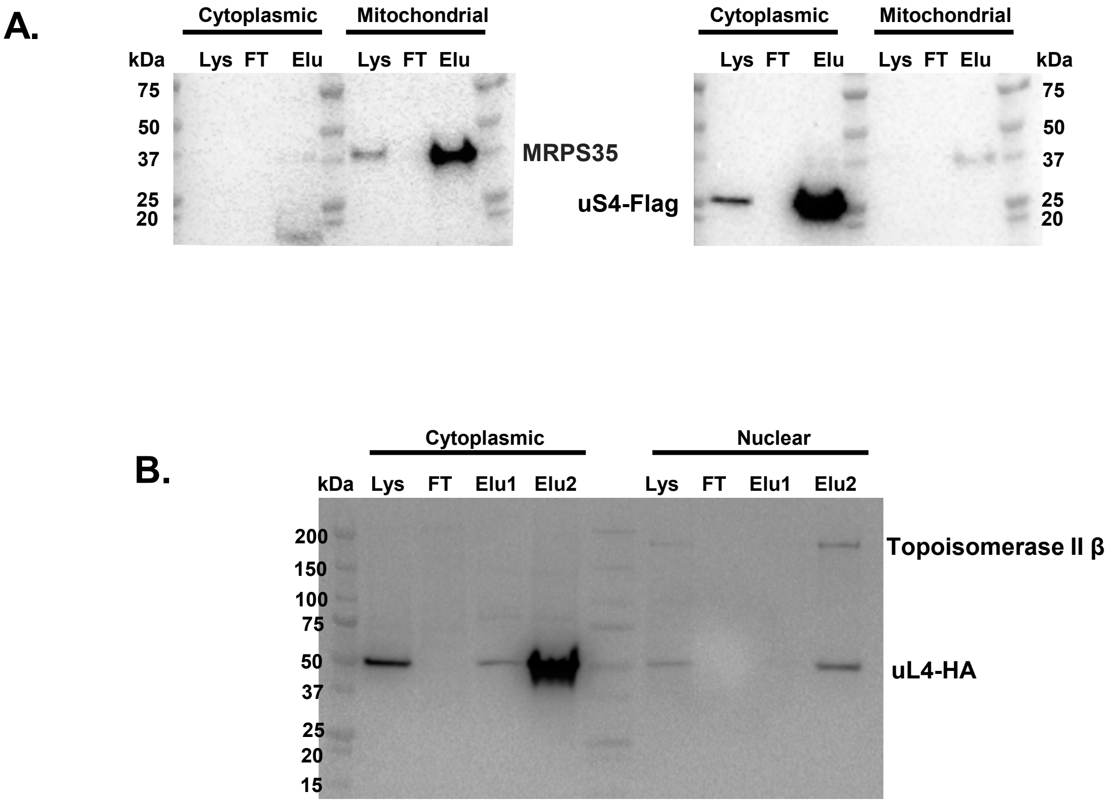


**Supplementary Figure 8. RAPPL can purify and enrich ribosomes from lysates of different organelles. A.** Western blot analysis using αMRPS35 and αFlag antibody on RAPPL eluates from mitochondrial and cytoplasmic fractions isolated from HEK-293 cells in which uS4 was Flag-tagged by CRISPR/Cas9. **B.** Western blot analysis using αTopoisomerase II β and αHA antibody on RAPPL eluates from cytoplasmic and nuclear fractions isolated from HEK-293 cells in which uL4 was HA-tagged by CRISPR/Cas9. Lysate (Lys), flow-through (FT), and RAPPL elution (Elu) are indicated in the figures Molecular weight markers indicate size of tagged uS-4. MRPS25, uL4 and Topoiosmerase II β protein.

### Supplementary Figure 9

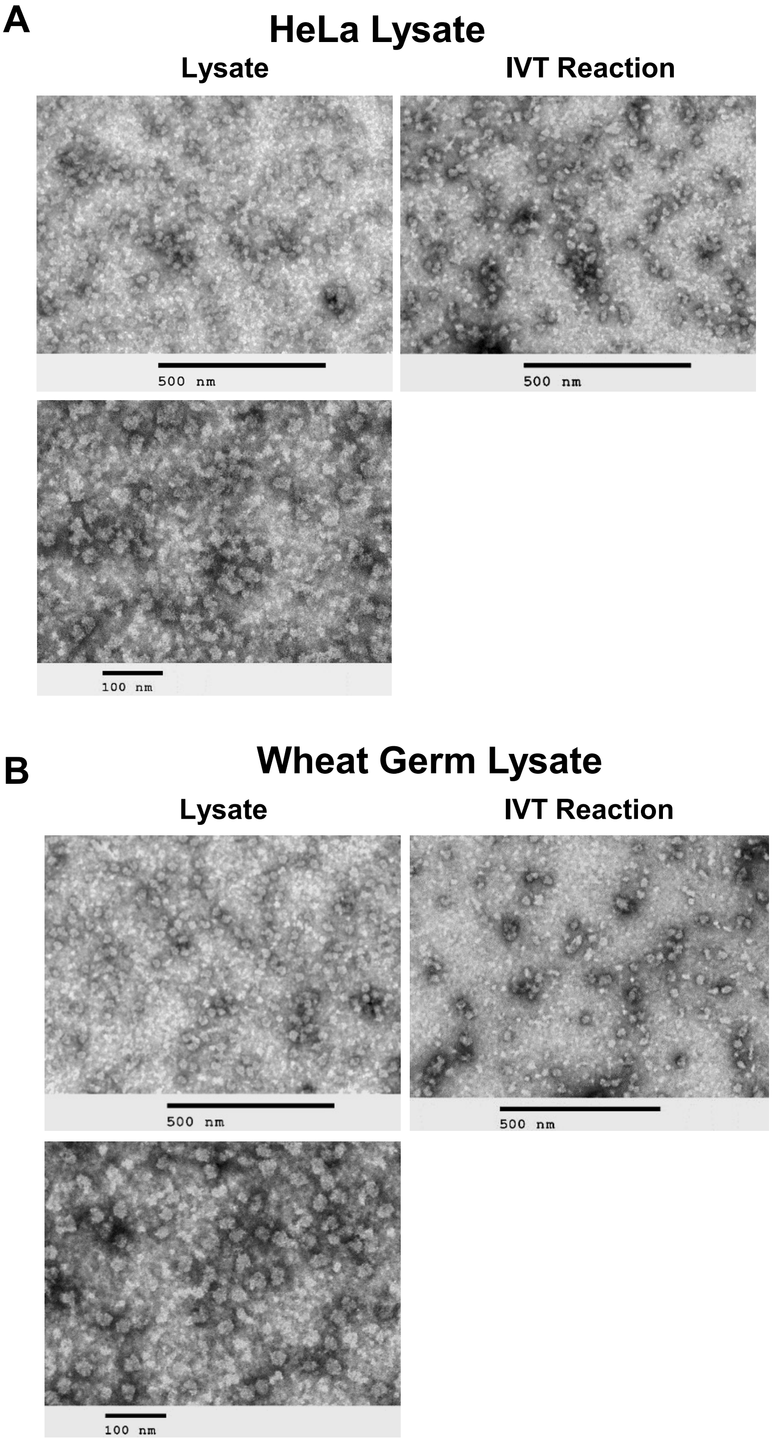


**Supplementary Figure 9. RAPPL can isolate non-translating and translating ribosomes from commercial cell lysates. A.** Thermo Scientific 1-Step IVT HeLa lysates in absence and presence of *in vitro* synthesized mCherry mRNA (IVT reaction). **B.** Promega TnT® Coupled Wheat Germ Lysate in absence and presence of *in vitro* synthesized mCherry mRNA (IVT reaction). All samples were subject to RAPPL and the eluates visualized by TEM. The scale bar represents 100 or 500 nm.

**Supplementary Figure 10**

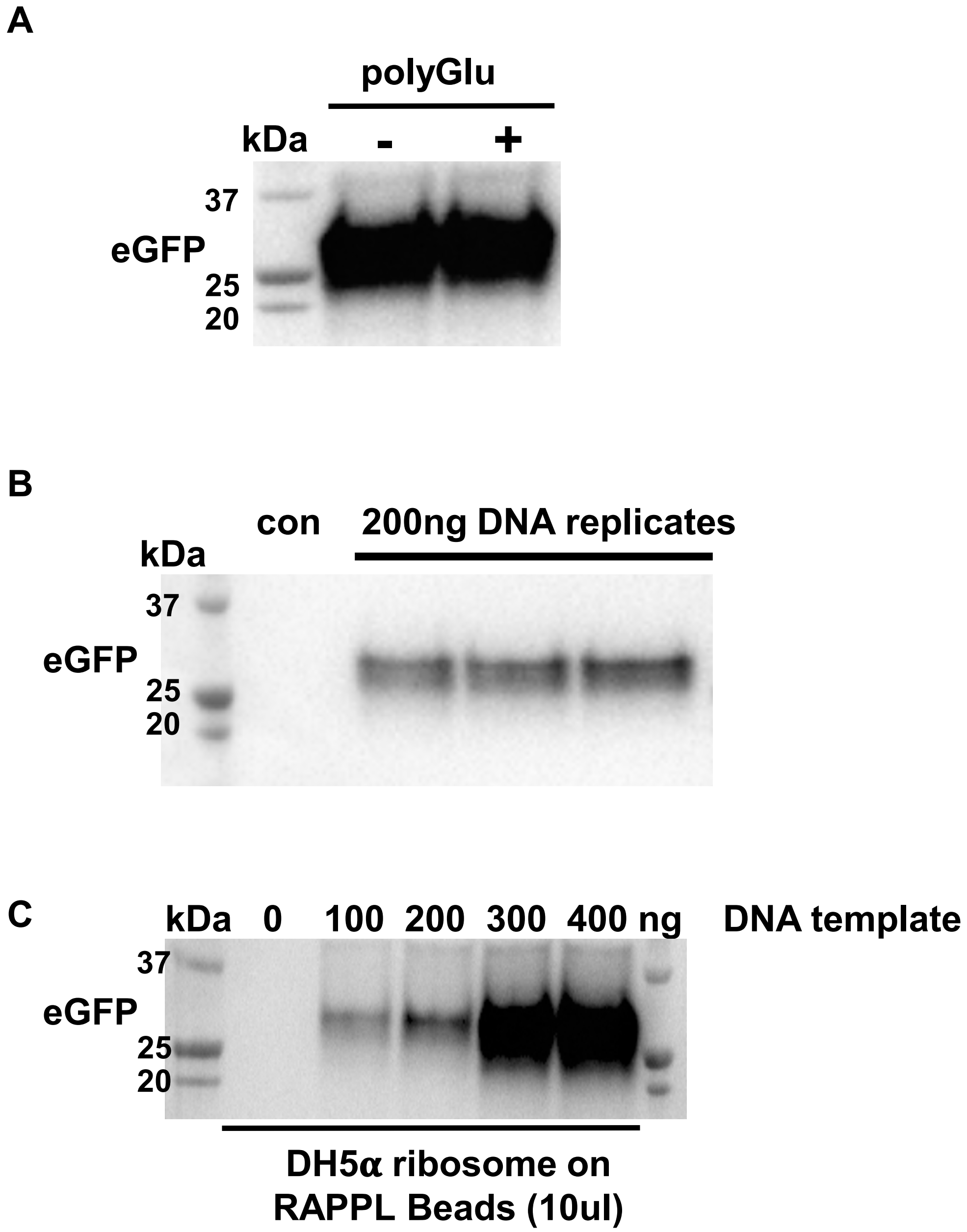


**Supplementary Figure 10. RAPPL purified *E. coli* ribosomes are translationally active. A.** Western blot analysis of NEB PURExpress *in vitro* reaction using eGFP DNA template. Reactions were executed in the absence (-) and presence (+) of poly-D-glutamate used in RAPPL elution buffer. **B.** Western blot analysis of eGFP protein synthesized in *in vitro* reactions using RAPPL isolated *E. coli Dh5α* ribosomes (1.5μg / μL). Western blot samples correspond to fluorescence reading shown in Figure 5B. **C.** PURExpress reactions in the presence of increasing amounts of eGFP DNA template and 10μL of poly-lysine (RAPPL) beads used to purify *E. coli Dh5α* ribosomes. αeGFP antibody was used to visualize synthesis of eGFP protein. Molecular weight markers indicate size of eGFP protein.

### Supplementary Figure 11

| STATE | Partially Rotated |
|---|---|
| Data collection statistics | |
| Microscope | Titan Krios G3 300kV Cryo-TEM |
| Camera | Gatan K2 Summit on GIF filter |
| Voltage (kV) | 300 |
| Magnification | 59kx |
| Total dose (e-/Å$^2$) | 55 |
| Defocus range (μm) | -0.6 to -2.0μm |
| Exposure Time (s) | 11.63 |
| Dose Rate (e/px/s) | 5.95 |
| Fractions | 50 |
| Calibrated pixel size (Å) | 1.122 |
| Micrographs collected | 2,498 |
| Initial number of particles | 646683 |
| Refined particles | 294300 |
| Symmetry | C1 |
| Map resolution (Å) | 2.7 |

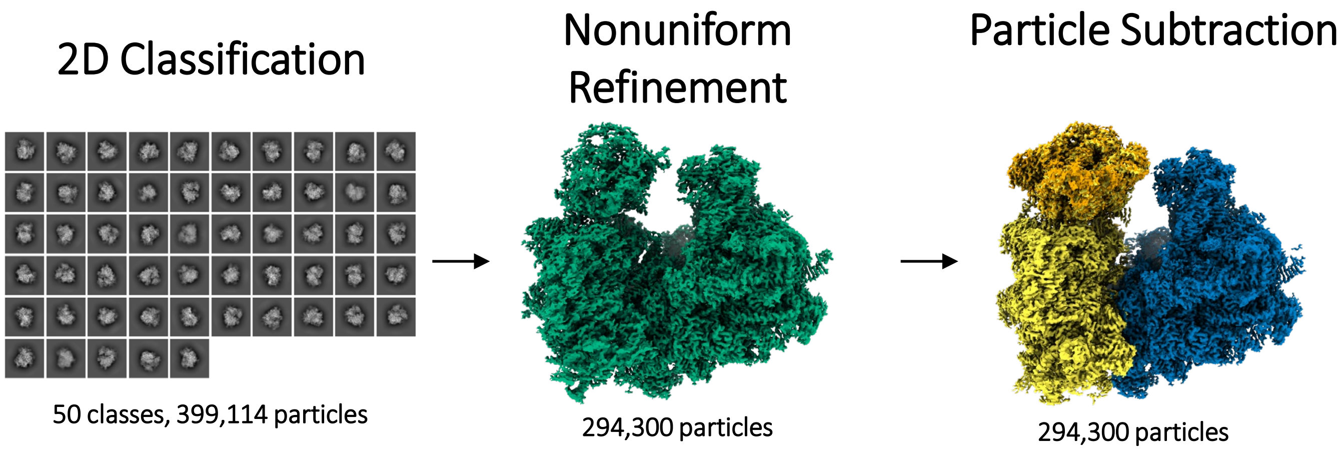


**Supplementary Figure 11. CryoEM processing of RAPPL purified and structurally characterized *C. neoformans* ribosomes.** Table representing cryoEM dataset collection parameters. Images indicate 2D classification, nonuniform refinement, and particle subtraction particle numbers.
